# In vivo spatiotemporal dynamics of astrocyte reactivity following neural electrode implantation

**DOI:** 10.1101/2022.07.01.498483

**Authors:** Sajishnu P Savya, Fan Li, Stephanie Lam, Steven M. Wellman, Kevin C. Stieger, Keying Chen, James R. Eles, Takashi D.Y. Kozai

**Affiliations:** Department of Bioengineering, University of Pittsburgh, Pittsburgh, PA, USA; Northwestern University, University of Pittsburgh, Pittsburgh, PA, USA; Center for Neural Basis of Cognition, University of Pittsburgh, Pittsburgh, PA, USA; Computational Modeling & Simulation PhD Program, University of Pittsburgh, Pittsburgh, PA, USA; Center for Neuroscience, University of Pittsburgh, University of Pittsburgh, Pittsburgh, PA, USA; McGowan Institute for Regenerative Medicine, University of Pittsburgh, Pittsburgh, PA, USA; NeuroTech Center, University of Pittsburgh Brain Institute, Pittsburgh, PA, USA

**Keywords:** Astrogliosis, foreign body response, intracortical microelectrode, neurovascular coupling

## Abstract

Brain computer interfaces (BCIs), including penetrating microelectrode arrays, enable both recording and stimulation of neuronal cells. However, device implantation inevitably causes injury to brain tissue and induces a foreign body response, leading to reduced recording performance and stimulation efficacy. Astrocytes in the healthy brain play multiple roles including regulating energy metabolism, homeostatic balance, transmission of neural signals, and neurovascular coupling. Following an insult to the brain, they are activated and observed to gather around the site of injury. These reactive astrocytes have been regarded as one of the main contributors to the formation of a glial scar which affects the performance of microelectrode arrays. This study investigates the dynamics of astrocytes within the first 2 weeks after implantation of an intracortical microelectrode into the mouse brain using two-photon microscopy. From our observation astrocytes are highly dynamic during this period, exhibiting patterns of process extension, soma migration, morphological activation, and device encapsulation that are spatiotemporally distinct from other glial cells, such as microglia or oligodendrocyte precursor cells. This detailed characterization of astrocyte reactivity will help to better understand the tissue response to intracortical devices and lead to the development of more effective intervention strategies to improve the functional performance neural interfacing technology.

## Introduction

Intracortical microelectrode arrays can measure neuronal information directly from the brain [1–4]. These neural interfacing technologies are used to understand neuroscience phenomena [5–11]. Additionally, knowledge gained from these devices can be applied clinically to treat neurological disabilities, such as restoring loss of motor or sensory function to tetraplegic patients [12–19]. However, limitations remain in the application of penetrating microelectrodes, such as large variability in long-term performance and stability [20–22]. The neuroinflammatory response that follows device implantation plays a significant role in biological and mechanical device failure modes [23–26]. Subsequent glial scarring and neuronal death results in a decrease in the number of neurons within the recording radius (80∼140 µm) [11, 27–30]. In turn, this leads to a drop in the number and quality of functioning electrodes over time [31].

Astrocytes are a major class of glial cell involved in tissue inflammation and scar formation within the central nervous system (CNS). They consist of 20% to 40% of all glia [32], and have long been a focus in microelectrode biocompatibility studies [33–35]. Under normal conditions, astrocytes have been described as tiling the entire CNS in a contiguous and non-overlapping manner with highly specialized [36, 37]. These cells perform a multitude of functions in homeostatic regulation of ions, neurotransmitters, water, and blood flow–all critical aspects of neural function [24, 38, 39]. Astrocytes can transmit intercellular Ca^2+^ waves in response to sensory processing and release glial transmitters in a Ca^2+^-dependent manner [40–42], thereby playing an active role in essential brain processes [43, 44]. However, these important homeostatic functions are disrupted following implantation of microelectrodes as astrocytes become reactive and facilitate the foreign body response (see review [24]).

Insertion of a microelectrode array not only inherently damages brain cells but also triggers several biological responses, including rupture of the blood-brain barrier and mechanical strain in the tissue [28, 45–47]. These responses initiate coordinated biological responses between several glial cell types such as astrocytes [48], microglia [49], and NG2 progenitor cells [50] (also known as oligodendrocyte precursor cells). The initial microglial response occurs within minutes of implantation and begins with the preferential process extension toward the site of injury [49, 51]. Recruited microglia participate in inflammation, the formation of giant cells, and phagocytosis of foreign debris [27, 52, 53]. Following implantation injury, microglia release cytokines which can activate nearby astrocytes, leading to process extension of astrocytes toward the injured area [54]. The switch to a reactive state is hallmarked by an astrocyte proliferation, pronounced upregulation of glial fibrillary acidic protein (GFAP), extension of processes toward the injured site, hypertrophy of soma and processes ultimately forming a glial scar [32, 33]. In turn, reactive astrocytes mutually coordinate with microglial via TNF-*α* and/or IFN-*γ* cytokines/chemokines [55, 56]. These astrocytes undergo proliferation and form scar tissue, termed astrogliosis, to prevent spread of secondary injury and mediate subsequential repair [24, 32, 33, 57].

While astrocytes have capacity of self-division, previous studies have demonstrated that astrocytes can also be generated from differentiation of recruited NG2 cells in the local parenchyma near implants [50, 58–60]. Astrocyte proliferation is essential for scar formation, reducing the spread and persistence of inflammatory cells, maintaining, and repairing the blood-brain-barrier (BBB), and limiting neuronal and non-neuronal tissue damage [24, 25, 60, 61]. For example, astrocytes play a role in producing glutathione and reducing reactive oxygen species, thus protecting neurons from oxidative stress [57, 62, 63] and neurodegenerative diseases [64, 65]. On the other hand, astrogliosis can be detrimental toward the injured CNS by impeding axonal and synaptic regeneration [66, 67]. Thus, understanding the role of astrocytes in neuroinflammation and foreign body response near the implanted device would contribute to a comprehensive picture of intrinsic biological reactions and future intervention strategies to mitigate implantation injury. Revealing the time course of morphological changes in astrocytes is one of the first step in identifying potential windows for therapeutics. However, this spatiotemporal mapping remains uncovered yet.

Previous immunohistochemistry studies have characterized glial responses in relation to chronic electrode implants [60, 68, 69], these endpoint histology studies are unable to capture real-time cellular dynamics. Here, we utilize two-photon microscopy to visualize astrocyte behavior and the spatiotemporal role of astrocytes in glial scar formation after insertion of a microelectrode *in vivo*. While previous two-photon studies demonstrated that astrocyte-related fluorescence and morphologies do not change from 2 to 10 weeks following probe implantation [48], post-mortem histology studies show significant changes in astrocytes following probe implantation compared to non-implant controls [30, 33, 52, 69–71]. Here, we show that the astrocyte response to microelectrode implantation is highly dynamic over the first two weeks post-implantation. Together with microglia [49], NG2 glia [50], as well as oligodendrocytes [72], astrocytes undergo remarkable morphological alternation, which may affect neuronal activity and the performance of the neural implant. Comprehensive characterizations of diverse glial subtype provide a complete picture of the integrated cellular responses during the foreign body response and provide various perspectives of intervention design to improve the stability and longevity of these functional intracortical devices.

## Methods

### Surgical Probe Implantation

Experimental methods were performed using previously established methods [31, 45, 49, 50, 53, 73–77]. Transgenic mice expressing green fluorescent protein (GFP) under two separate astrocyte-specific promoters were used (Tg(ALDH1L1-GFP)D8Rth/J, n = 5 and Tg(GFAP-GFP)14Mes/J, n = 5), as well as microglia (CX3CR1-GFP/J, n=5) and NG2 glia (Tg(Cspg4-GFP)HDbe/J, n = 5) (Jackson Labs, Bar Harbor, ME). Acute studies (0 -6 h) were carried out using ALDH1L1-GFP animals due to the preferential labeling of the astrocytic processes and ability to visualize endfeet at acute time points. GFAP-GFP were used for chronic imaging studies (1 h – 14 d) due to increase in fluorescence of astrocytes during the chronic foreign body response. Microglia and NG2 glia were imaged from 0-72 hrs as previously published [49, 50]. Mice were intraperitoneally (IP) injected with a mixture of 75 mg/kg of ketamine and 7 mg/kg of xylazine (IP) to induce anesthesia and fixed onto a stereotaxic surgical frame. Anesthetic updates were administered using 40 mg/kg of ketamine. The skin atop of the skull was cleaned with anti-septic betadine followed by ethanol prior to skin removal. Connective tissue was removed, and the surface of the skull was dried prior to application of 1-2 drops of Vetbond. A single bone screw hole was drilled over each motor cortices using a high-speed dental drill. Bone screws were inserted into each hole at a 45° angle into the caudal direction and stabilized using dental cement. Over the left visual cortex – center point 1.5 mm rostral to lambda and 1 mm lateral to midline – a 4mm by 6mm craniotomy was formed. During drilling, saline was applied regularly to avoid thermal damage. Once the skull was removed, a multi-shank Michigan microelectrode array (A4×4-3mm-50-703-CM16; NeuroNexus Technologies, LLC, Ann Arbor, MI) was inserted at a 30° angle at a speed of ∼200 μm/s into the cortex in the rostral direction parallel to midline for 600 μm (400μm in horizontal plane) using an oil hydraulic Microdrive (MO-82g, Narishige, Japan). With these insertion parameters, the tips of the electrode are positioned at about 300 μm below the surface of the brain (Fig. 1a). Blood vessels were avoided as much as possible during insertion. Animals intended for acute insertion studies were implanted under the two-photon microscope. For chronic imaging, a cranial window was installed by using Kwik-Sil to seal the craniotomy prior to placing a glass coverslip over the probe. The edges of the coverslip were then secured and sealed with dental cement. A well was made around the imaging window between 1.5mm to 2mm tall to hold saline for imaging via a 16x water-immersive 3.5mm working distance objective lens (Nikon Instruments, Melville, NY). All procedures and experimental protocols were approved by the University of Pittsburgh, Division of Laboratory Animal Resources, and Institutional Animal Care and Use Committee in accordance with the standards for humane animal care as set by the Animal Welfare Act and the National Institutes of Health Guide for the Care and Use of Laboratory Animals.

**Fig. 1.**
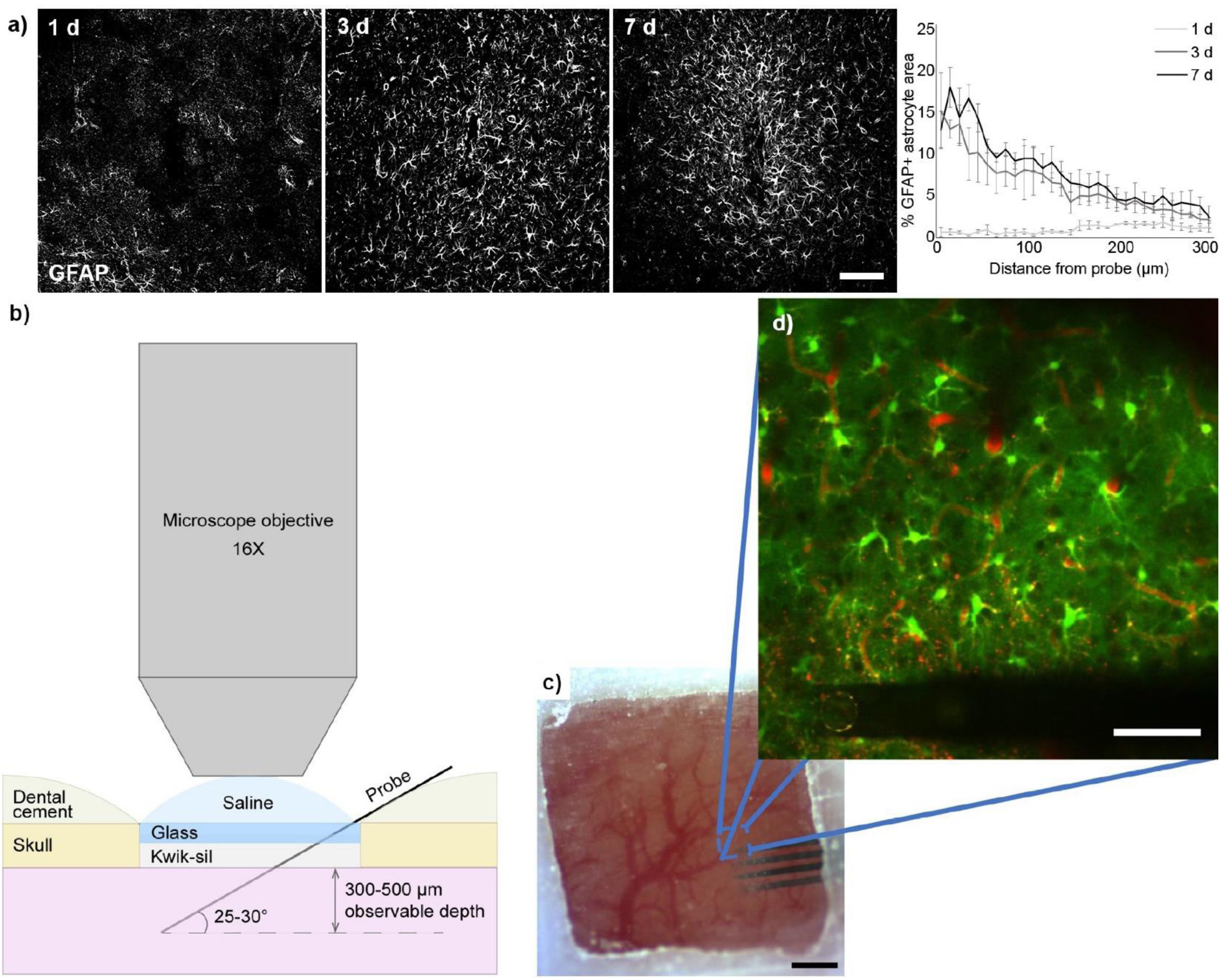
Experimental set-up to track astrocyte dynamics following device insertion. a) Immunolabeling for GFAP in implanted tissue (*n* = 3) demonstrates elevation in GFAP+ astrocytic area within first week following insertion. Scale bar = 100 μm. b) Schematic of chronic imaging window for *in vivo* visualization of implanted microelectrodes using two-photon microscopy. The probe was implanted at a 30° angle within the cortex and sealed with Kwik-Sil prior to placing a cover glass on top and securing the window with dental cement. c) Surgical image of cranial imaging window. Scale bar = 1 mm. d) Inset of (c). Astrocytes (*GFAP-GFP* or *ALDH1L1-GFP*) were quantified and observed up to 400 μm from the probe surface. Scale bar = 50 µm.

### Two-photon Imaging

For longitudinal imaging, mice were anesthetized using isoflurane (1.0-1.5%, mixed with 0.9 L/min O_2_). To visualize blood vessels, mice were injected with 0.2 mL of sulforhodamine 101 (SR101) preceding every Z-stack taken or every hour. Mice exhibiting pial surface bleeding which resulted in poor imaging quality were removed from the study. Probe shanks and adjacent regions of tissue were imaged using a 412.8 x 412.8 μm (1024 x 1024 pixels) field of view. The first and last shank with the least amount of surface vasculature were used to determine the imaging region for maximum image clarity. Z-stack images spanned the entire depth of the probe implant (300 μm) with a 2 μm step-size and ∼5s/frame scan rate and were taken every hour up to 12 h post-insertion, as well as every 24 h up to 14 d post-insertion.

### Data Analysis

Analysis of putative astrogenesis used z-stacks taken at 0.25 d, 0.5 d, 1 d, 3 d, 4 d, 7 d, 10 d, and 14 d post-insertion. To obtain the number of new cells per animal, ImageJ was used to generate z-projections by averaging the intensities of the z-stacks every 10 images (20 µm depth). Six z-projections (120 µm depth in total) were used per animal. Z-projections were sorted to match the imaging depth of the z-projections at the current and previous time points. Next, z-projections from the current time point were aligned to the previous time point’s z-projections using ‘TurboReg’ plugin [78] to perform a rigid body transformation. Following alignment, z-projections of the same plane in the previous and current time points were compared by making a composite image of the aligned z-projections. Additionally, z-projections were concatenated into a single stack to confirm new cells by alternating between the two images for comparison. Cells not observed in the previous time point’s z-projections were verified in the original z-stack to confirm that the cell was not drifting between time points. Once confirmed that the cell is new, cells were classified as proliferating from either dividing or differentiating cells by observing the new cell’s surroundings. Cells were classified by the contact of their cell body with surrounding cell bodies, where no contact and contact corresponded with differentiation and division, respectively. The number of new cells per day was calculated by dividing the number of new cells within the entire analyzed volume (∼12 µm^3^) over the elapsed time between consecutive time points. The average number of new cells per day was calculated by averaging the number of new cells per day for all animals at each time point. Number of new cells per mm^3^ was obtained by dividing the number of new cells between subsequent time points in all of the six z-projections by analyzed area. The analyzed area excluded the areas in the image that were covered by the probe, had no fluorescent signal, and were unaligned between consecutive time points.

ImageJ was used to process acquired z-stack images. Z-stacks of consecutive time points were aligned using the probe as a reference point to quantify astrocyte process extension/retraction, soma diameter, soma migration, and morphology. In all cases the control was the first image immediately following insertion to in order to account for insertion drag related tissue shifts. TurboReg was used to calculate offsets in one z-stack in relation to another. Subsequently, the z-stacks were aligned using the translate function. The movement of processes and soma surrounding the probe surface were quantified from onset of movement to point of destination. Movements were tracked using the “measure” feature in ImageJ and noting the XY coordinates of the center point of tips of processes or center of soma between consecutive imaging time points. Directionality of processes and migration of soma were determined by hemisecting the soma such that the hemispheric divide is parallel to the probe’s surface. Process velocity and soma velocity recordings were defined as the distance between consecutive time points of process end-feet and soma center point, respectively. Positive direction indicated movement towards the probe and negative direction indicated movement away from the probe. Therefore, if movements in the positive direction and negative direction were equal, the plot would be centered on zero with some standard deviation. Quantification of process distance and soma distance from the probe was done by first drawing a radius around the shanks. The radius was created by drawing a best fit curve of the astrocytic processes closest to the shanks. Similarly, another radius was created using the soma closest to the shanks. Distances between the processes oriented toward the shanks or the soma closest to the shanks (formulated radius) and the surface of the shank was measured. Peak velocity was determined by isolating the largest process and soma velocity over the course of 3 d. Activation of astrocytes was defined as the point of process extension past baseline threshold. Each astrocyte was analyzed to see whether or not they exceeded this threshold. Astrocytes with processes extended beyond the threshold were denoted as activated.

Coverage of astrocyte processes on the surface of the probe was quantified as the measured fluorescence over the total probe area. Using the “Interactive Stack Rotation” feature of ImageJ, z-stacks were rotated to re-slice the total volume of the tissue normal to the surface of the probe. By utilizing an ImageJ thresholding method that utilizes an isodata algorithm, a binary mask of the slice projection of a small volume of tissue immediately above the surface of the probe was made [79]. The probe was outlined and the quantity of non-zero pixels within the outline was measured and taken as a fraction of the total area. The percent change in astrocyte cell body area was calculated as the difference between cell body area at the current time point and cell body area at 1 h divided by the cell body area at 1h post-insertion time point.

Astrocyte, microglia, and NG2 glia morphologies were characterized using metrics previously outlined [77]. Briefly, astrocytes were classified as either ramified (non-activated) or activated (transitional stage), denoted as 1 or 0, respectively. Astrocytes were binned in 50μm increments away from the probe surface. As previously outlined [77], a logistic regression was used to fit the data to show the distribution of astrocytes in a ramified or activated state as a function of distance from the probe surface. A transitional index (T-index) and directionality index (D-index) were used to quantify changes in morphology as well. T-index was determined by measuring the length of the longest process extending toward the probe (n) and the length of the longest process extending away from the probe (f). D-index was determined by denoting a hemispheric divide of each astrocyte and using this divide to distinguish between the number of processing extending toward the probe (n) and the number of processes positioned away from the probe (f). The formula used to calculate index values is as follows:

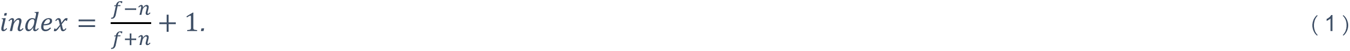

As with the ramification index, an index of 1 denotes a ramified state, while an index of 0 denotes a fully activated state for both T- and D-index. Astrocytes were binned in 50 μm increments away from the probe surface. A dual sigmoidal function was used to fit the index distribution, as previously detailed [50].

Vasculature changes were quantified by measuring blood vessel diameter over a z-stack projection of the entire blood vessel volume. Percent change was calculated by determining the difference of diameter between the current time point and the 1 h time point and dividing by the 1 h post-insertion time point. Astrocyte soma volume was quantified by outlining the soma of individual astrocytes using ImageJ.

#### Immunohistochemistry

For Immunohistochemistry, implantation of microelectrode arrays were performed as described previously [28]. Prior to surgery, all electrode arrays and surgical tools were sterilized using ethylene oxide for 12 hours. Nonfunctional single shank Michigan-style microelectrodes (A16-3 mm-100-703-CM15) were perpendicularly implanted into the left primary monocular visual cortex (V1m) of 8-week old C57BL/6J male mice (Jackson Laboratory, Bar Harbor, ME). Anesthesia and surgical preparation was conducted as describe above. For these mice, three bone screws were drilled into the bone over both motor cortices and the contralateral visual cortex to help secure the headcap. A 1 mm drill-sized craniotomy was formed at 1 mm anterior to lambda and 1.5 mm lateral from the midline using a high-speed dental drill with periodically administered saline to prevent overheating of the brain surface. Gelfoam was used to prevent the brain from drying out prior to implantation. The microelectrode array was carefully inserted using a stereotaxic manipulator at a speed of ∼2 mm/s until the last electrode contact site was below the surface (∼1600 μm from the pial surface). The inserted probe was sealed using a silicone elastomer (Kwik-sil) and a headcap was secured using UV-curable dental cement (Henry Schein, Melville, NY). Ketofen was administered at 5 mg/kg up to 2 days post-operatively.

Mice were sacrificed at 1, 3, and 7 d post-insertion (*n* = 3 per time point) for immunohistochemical analysis in MATLAB as described previously [60]. Mice were transcardially perfused with 1x PBS followed by 4% paraformaldehyde prior to decapitation and post-fixation at 4°C overnight. Brains were then extracted and sequentially soaked in 15% and then 30% sucrose at 4°C for 24 hrs each. After the samples had reached equilibrium they were frozen in 2:1 20% sucrose in PBS:optimal cutting temperature compound (Tissue-Tek, Miles Inc., Elkhart, IN, United States). The samples were then sectioned horizontally (25 μm slice thickness) and mounted onto glass slides using a cryostat (Leica Biosystems, Wetzlar, Germany). Sections were re-hydrated with 1x PBS washes for 30 min prior to staining. For antigen retrieval, sections were incubated in 0.01 M sodium citrate buffer for 30 min at 60°C followed by incubation in a peroxidase blocking solution (10% v/v methanol and 3% v/v hydrogen peroxide) for 20 min at RT. To permeabilize cells, the sections were then incubated in 1% triton X-100 with 10% donkey serum in PBS for 30 min. Excess detergent was rinsed off with 1x PBS (8×4 min) washes. Then, sections were incubated with the following primary antibodies: chicken anti-GFAP (Abcam cat. AB4674, 1:500), mouse anti-Ki67 (BD biosciences cat. 550609, 1:50), and mouse anti-Olig2 (Sigma cat. MABN50, 1:100) overnight at 4°C. Tissue sections were then washed in 1x PBS (3×5 min) before incubation with donkey anti-chicken IgY 647 (Sigma-Aldrich, St. Louis, MO, United States) and donkey anti-mouse IgG 488 (Thermo Fisher Scientific, Waltham, MA, United States) at 1:500 dilution in 1x PBS for 2 h at RT. Sections were then coversipped using Fluoromount-G (Southern Biotech, Birmingham, AL, United States). Labeled samples were imaged using a confocal microscope (FluoView 1000, Olympus, Inc., Tokyo, Japan) with a 20x oil-immersive objective lens.

### Statistical Analysis

Statistical significances of process extension and soma migration were determined using a one-way ANOVA test. A repeated measures ANOVA was used for data comparisons within astrocyte populations. Dunnett’s post-hoc test with a Bonferroni correction was used to compare T-index and D-index of 10 min against all other time points at each bin. Morphological changes between astrocytes, NG2 glia, and microglia were determined using a two-way ANOVA test with Holm-Sidak post-hoc test. An independent variable t-test and Pearson correlation coefficient were used to compare data between vasculature and astrocytes. A *p*-value of *p*<0.05 was used to show significant differences.

Analysis of differences between newly generated cells from differentiation and division were done at each time period using Wilcoxon’s matched pairs signed rank test. Differences in the effect of time on the rate of newly generated cells at each time period was tested with repeated measures Friedman test (nonparametric) with Dunn’s post-hoc test with Bonferroni adjustment to compare 0.25 d (control) to all other time points. A *p*-value of *p* < 0.05 was considered significant. To analyze the effects of distance and time on the new cell density between time periods, a repeated-measures two-way Friedman test with Dunn’s test and Bonferroni adjustment was used.

## Results

Although it is well known that astrocyte density and GFAP intensity increases around chronically implanted microelectrode arrays [33–35], it remains unclear how and when these increases occur [48]. SNR thresholded pixel count for GFAP+ pixels demonstrate an increase of GFAP+ area (Fig. 1a), demonstrating that the nature of GFAP intensity is not simply due to the upregulation of the GFAP marker, but also an increase in which the astrocytes occupy tissue volume. In order to characterize the acute (0 - 6 h) and chronic (1 h – 14 d) astrocyte response to microelectrode insertion, two mouse models expressing GFP in astrocyte through either the Aldh1l1 or GFAP promotor was used (Fig. 1). Aldh1l1-GFP mice display strong GFP labeling of astrocyte processes and endfeet, and were therefore used for quantifying process dynamics during the acute phase of the astrocyte response (0 – 6 h). GFAP expression was expected to increase during the chronic period of the astrocyte response, therefore GFAP-GFP mice were used to quantify the astrocyte dynamics during the chronic time points (1 h – 14 d). Prior to longitudinal imaging experiments, immunohistochemical labeling for GFAP revealed astrocyte reactivity, hypertrophy, gliosis, and overall increased area occupied within 300 μm from the probe surface within the first 7 d post-insertion (Fig. 1a). To uncover the dynamic spatiotemporal behavior of individual reactive astrocyte cells within the electrode-tissue interface, microelectrodes were implanted within transgenic animals with GFP-labeled astrocytes underneath a cranial window to allow optical access using two-photon microscopy (Fig. 1b). Astrocytes were found to have no morphological changes due to the craniotomy prior to probe insertion. Surface vasculature was avoided during insertion to limit bleeding. The laser power was kept at ∼20 mW (never exceeding 40 mW) [53] to prevent thermal damage and photomultiplier tube (PMT) settings were tuned for best image quality. A 300 µm region adjacent to the probe shank was imaged for quantification (Fig. 1c, d).

### Astrocytes initiate process extension, become hypertrophic and polarize towards the implanted electrode within 48 h

Astrocyte reactivity to CNS injury is heterogeneous can involve changes such as process extension toward the site of injury, hypertrophy, proliferation and glial scar formation, as well as substantial molecular changes [80–82]. Therefore, we first quantified the temporal progression of astrocyte process polarization and extension toward the electrode (Fig. 2). Briefly, process extension towards the probe was positive, while process extraction away from the probe was negative. In non-implanted controls, the average velocities were 0. Process extension velocity peaked at a velocity of 0.24 ± 0.15 µm/h 30 min after electrode insertion then quickly decreased within the next hour. Interestingly, process extension increased again after 6 h moving at a peak rate of 0.15 ± 0.02 µm/h. Process velocity stabilized between 4-7 d post implant; therefore, quantification is only included up to 7 d. Astrocytes within 50 µm of the probe surface extended their processes toward the probe faster than astrocytes farther away (Fig. 2c), suggesting that astrocytes closer to a site of injury may respond more to the injury due to a gradient of ATP from the injury site, astrocyte processes within 20 µm of the electrode, and particularly the processes facing the electrode were the first to begin extension (30 min; data not shown).

**Fig. 2.**
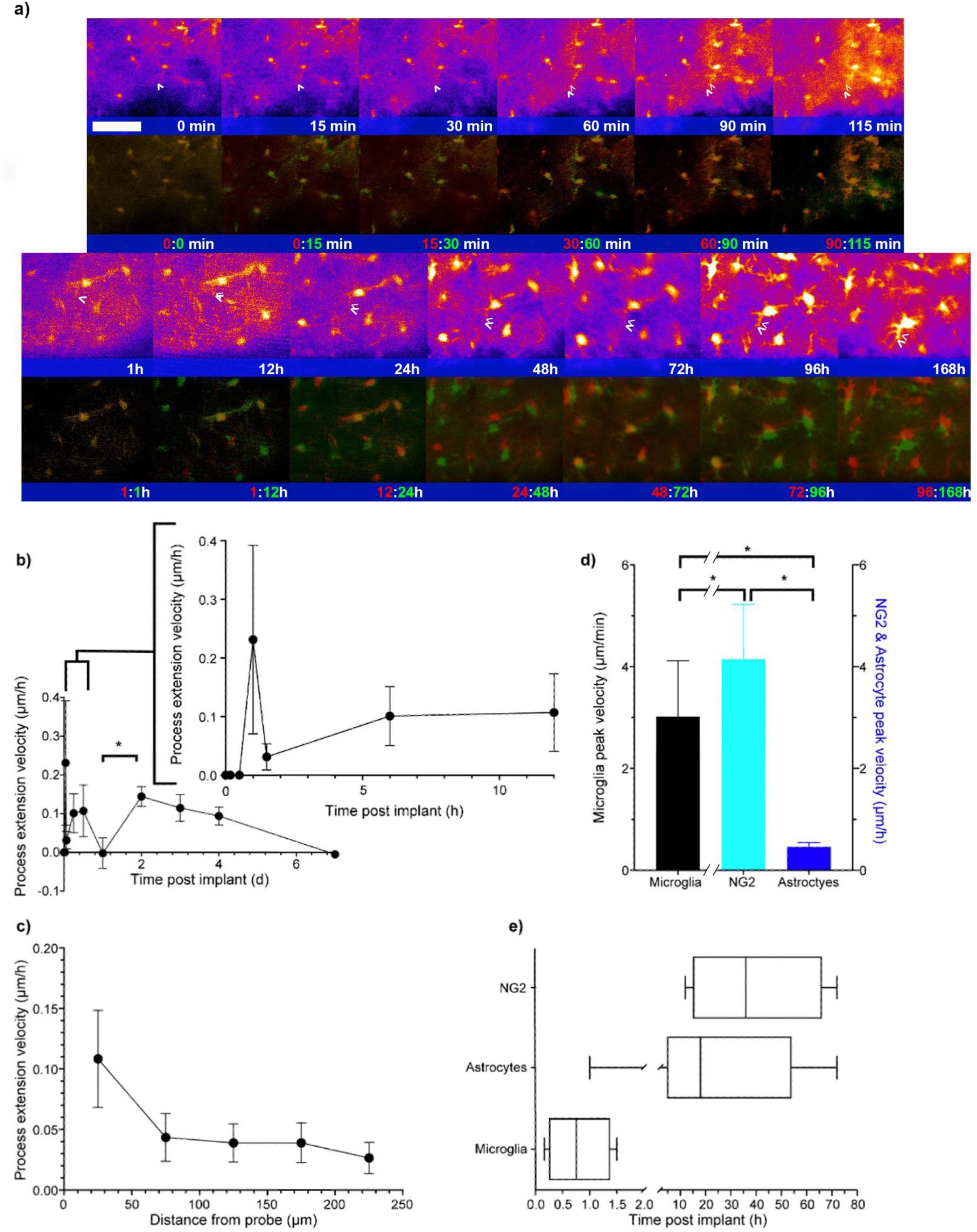
Astrocyte processes start to extend toward the probe at 1 h post-insertion. A) Superimposed images depicting astrocytes extending their processes toward the probe surface (*shaded blue*). Images of 20 µm z-projections centered on the processes are shown in ‘fire color’ on the top row. Two consecutive time points are merged into two color channels: red denotes the earlier time point and green denotes the later time point. Yellow depicts overlap between two consecutive time points. White arrows indicate the examples of process extension. Scale bar = 15 µm. b) Astrocyte process extension velocities were tracked over time. C) Rate of process extension versus distance from the probe surface demonstrates an increase in extension velocity for astrocyte processes closer to the device. D) Peak extension velocities reveal differences in process dynamics between microglia, NG2 glia, and astrocyte populations. Note the units for peak extension velocity differ for microglia cells (μm/min) compared to NG2 glia and astrocytes (μm/h). E) Distribution of the initiation of process migration of microglia, astrocytes, and NG2 glia reveal that astrocytes facilitate process extension in the period between microglia and NG2 glia populations. Information about microglia and NG2 data can be found in the footnote ^1^. * indicates p < 0.05.

While astrocytes are a critical component of the foreign body response to injury that have been suggested to contribute to degradation in device performance, microglia and NG2 glia have also been shown to process polarization toward the implant site. Our group has previously quantified the response of microglia and NG2 glia to electrode implantation up to 72 hours post insertion. Both microglia [83] and NG2 glia [57] are known to have coordinated communication and interaction with astrocytes during the foreign body response. Therefore, in order to provide a more comprehensive view of our current understanding of the acute-to-chronic glial response we compared the time course of astrocyte responses (0 – 14 d) with that of microglia and NG2 glia (0 - 3 d). In comparison, peak microglia velocities were significantly different from peak process velocities of NG2 glia, and peak astrocyte process velocities were significantly different from both peak process velocities of microglia and NG2 glia (Fig. 2d; *p* < 0.05). Interestingly, astrocyte processes moved at a rate of microns per hour, similar to NG2 glia. However, astrocytes were still considerably slower than NG2 glia [50]. While a limited number of astrocytes responded immediately after microglia, the bulk of astrocytes activated before NG2 glia around 20 h following electrode insertion (Fig. 2e).

While there was measurable movement of astrocyte processes towards the injury, it was clear that astrocyte reactivity was not limited to process movement. Hypertrophy of astrocytic processes and cell bodies follows injury and signifies astroglial reactivity (see reviews [32, 57]). Although upregulation of GFAP expression does not necessarily indicate reactivity, it does correlate with cellular hypertrophy in injury [84] and is understood to contribute to the modulation of process reorganization and motility [85]. Therefore, we next quantified astrocyte hypertrophy as the change in soma area between subsequent time points (Fig. 3). The area of astrocyte soma significantly increased 48 h after electrode implantation followed by a subsequent plateau (Fig. 3a-b; p<0.001). By 48 h post insertion, astrocyte soma area increased by 66 ± 9%, compared to 1h post insertion and remained enlarged throughout the 14-day study possibly indicating persistent reactivity. Interestingly, while astrocyte soma and process movement was elevated within the first 50 µm, the soma between 100 and 150 µm demonstrated the greatest hypertrophy (Fig. 3c).

**Fig. 3.**
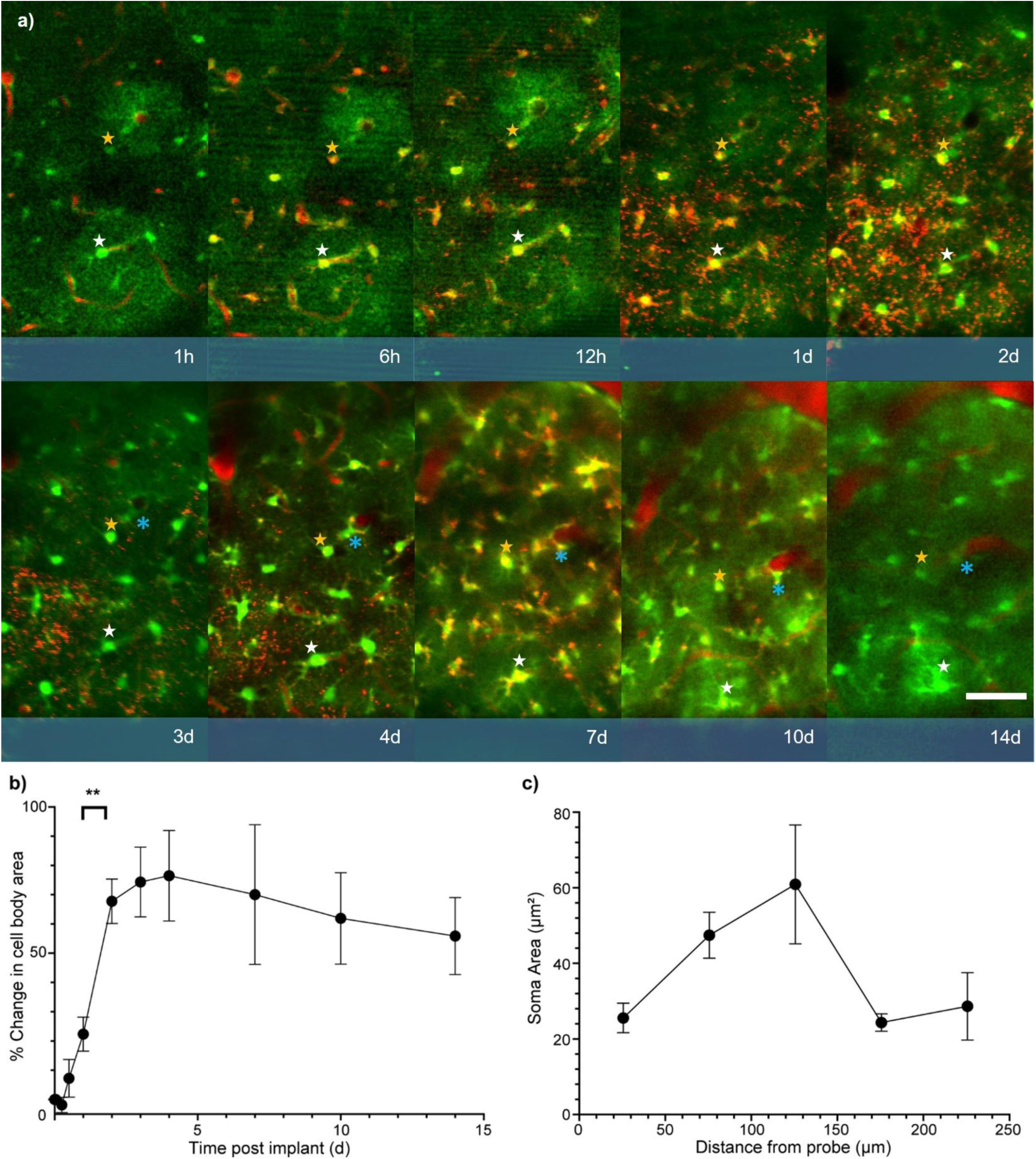
Astrocyte soma area increases over first initial days following implantation. A) Representative images of 20 µm thick z-projection show ramified or activated astrocytes (green) over 14 d with respect to the probe surface (*shaded blue*). Vasculature was labeled using SR101 (red) Asterisks indicate examples of measured soma. Scale bar = 50 µm. b) Percent change in soma area over time with respect to soma area at 1 h post-insertion reveals a significant increase in astrocyte soma area (hypertrophy) up to 3-5 d post-insertion. C) Astrocyte soma area plotted as a function of distance from the probe reveals a preferential increase in soma area within 50 to 150 µm from the probe surface. * indicates p < 0.05, ** p < 0.001.

A recent report uncovered that 99.8% of gray matter astrocytes contact at least one blood vessel [86]. It was suggested that this ubiquitous connection between astrocytes and blood vessels acts as a redundant mechanism to redistribute metabolites in conjunction with gap junction coupling in certain physiological states (e.g. pathological conditions) [86, 87]. Additionally, we have previously demonstrated that pial blood vessel diameter increases in the first 3 days following electrode insertion [50]. Therefore, we quantified whether astrocyte hypertrophy is correlated to changes in blood vessel diameter (Fig. 4). Blood vessels near the implanted probe increased in diameter over 14 d post-insertion (Fig. 4a). Quantifying the percent change in blood vessel diameter over time (taken as the difference in lumen diameter with respect to the diameter measured at 1 h post-insertion) demonstrated an increase in diameter at 6 h following probe insertion and a significant decrease at 12 h (Fig. 4b, *p*<0.05). Similar to the previous report [50], the diameter increases significantly during 2 to 3 d from 28 ± 7% to 75 ± 16% (Fig. 4b, *p*<0.05). Interestingly, this increase in vascular diameter was moderately correlated with the percent change increase in astrocyte soma area (Fig. 4c, R^2^=0.5586), suggesting a potential coupling between astrocyte behavior and vascular reactivity to device implantation.

**Fig. 4.**
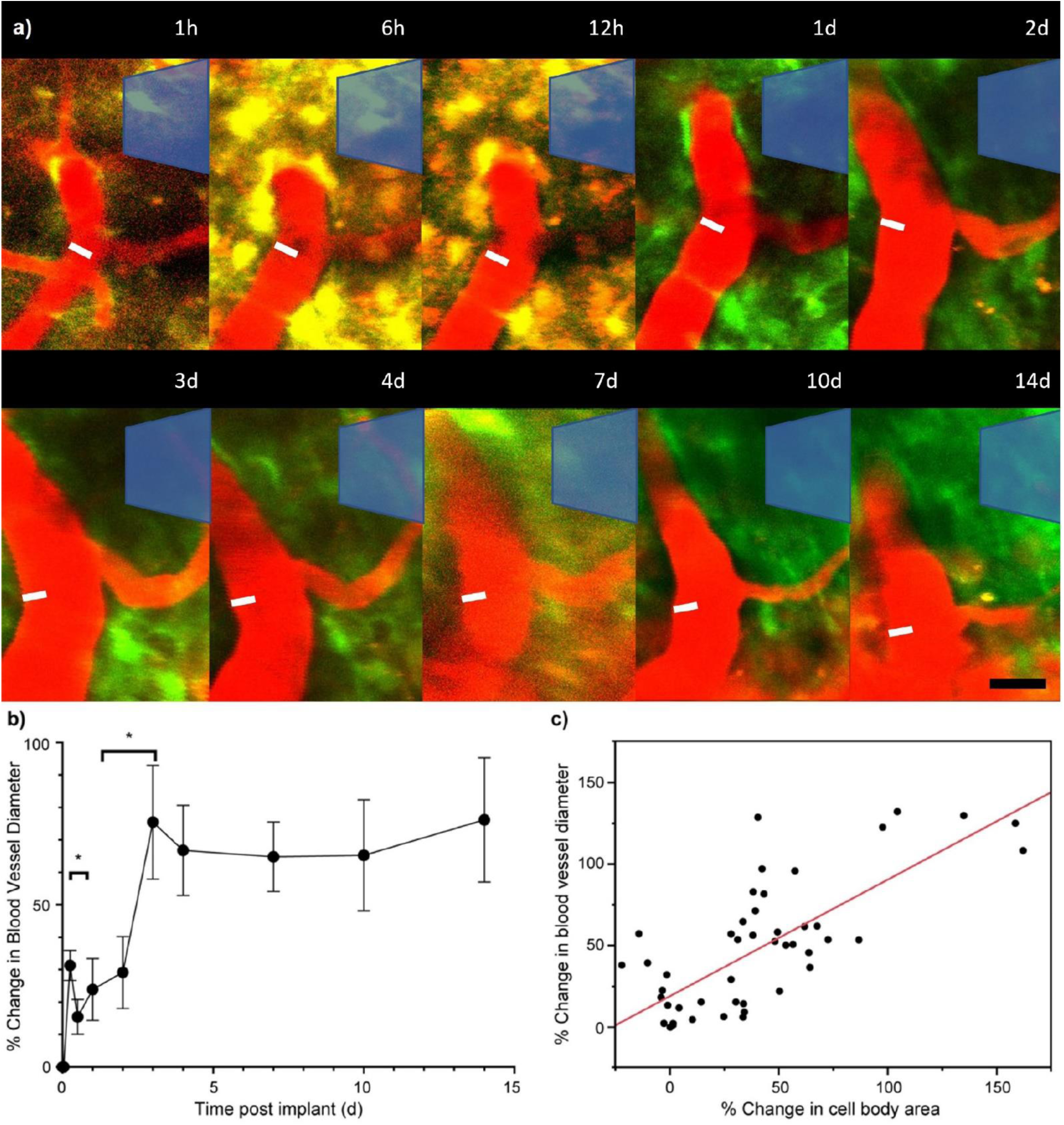
Dynamic changes to local vasculature following probe implantation. a) Images show 20 µm thick z-projection centered around the widest portion of blood vessels. Blood vessels adjacent to the probe (*shaded blue*) increased in diameter over 14 d following probe insertion. White line denotes the size of blood vessel at 1 h post-implantation. Note that GFP+ astrocytes appear yellow within initial 24 h following implantation due to cellular uptake of SR101 dye (*red*). Black scale bar = 25 µm. b) Percent change in vessel diameter compared to the diameter measured at 1h post-insertion. Blood vessels appear to significantly expand in diameter (dilation) up to 3 d post-insertion before stabilizing. c) Relationship between vessel diameter and the area of associated astrocyte soma reveal a positive correlation between astrocyte hypertrophy and vascular dynamics following device implantation. Data are collected from all time points in (a). * indicates p < 0.05.

Now that we have demonstrated that astrocytes extend processes towards the probe and increase soma area, we examined the extent of activation as a function of distance over time. During CNS injury and astrocyte reactivity, astrocyte upregulation of GFAP can aid the transition from ramified morphology to polarization in the direction of the injury site depending on the severity of the injury. Therefore, astrocytes were classified as either ramified (R) or transitional (T) by observing the orientation of their processes after bisecting the soma with an imaginary line parallel to the probe surface (Fig. 5a). Although a significant increase in hypertrophy did not occur until 48 h after insertion, astrocytes within 100 µm of the probe surface began to transition into a reactive morphology 1 h after insertion. In areas further than 100 µm of the probe surface, astrocytes activated much later (at 3 d post-insertion) (Fig. 5b). Analysis of the leading and lagging astrocyte processes in relation to the probe, quantified as T-index, showed astrocytes retracting processes which were oriented away and extending processes which were oriented toward the probe (Fig. 5c). There was greater preference for process extension toward the probe starting at 1 h after insertion for astrocytes within the first 200 µm (Fig. 5c). In addition, D-index quantification showed the number of processes facing toward the probe gradually increased starting at 6 h post-insertion for astrocytes within the first 100 µm and 7 d after insertion for astrocytes beyond 100 µm from the probe surface (Fig. 5d).

**Fig. 5.**
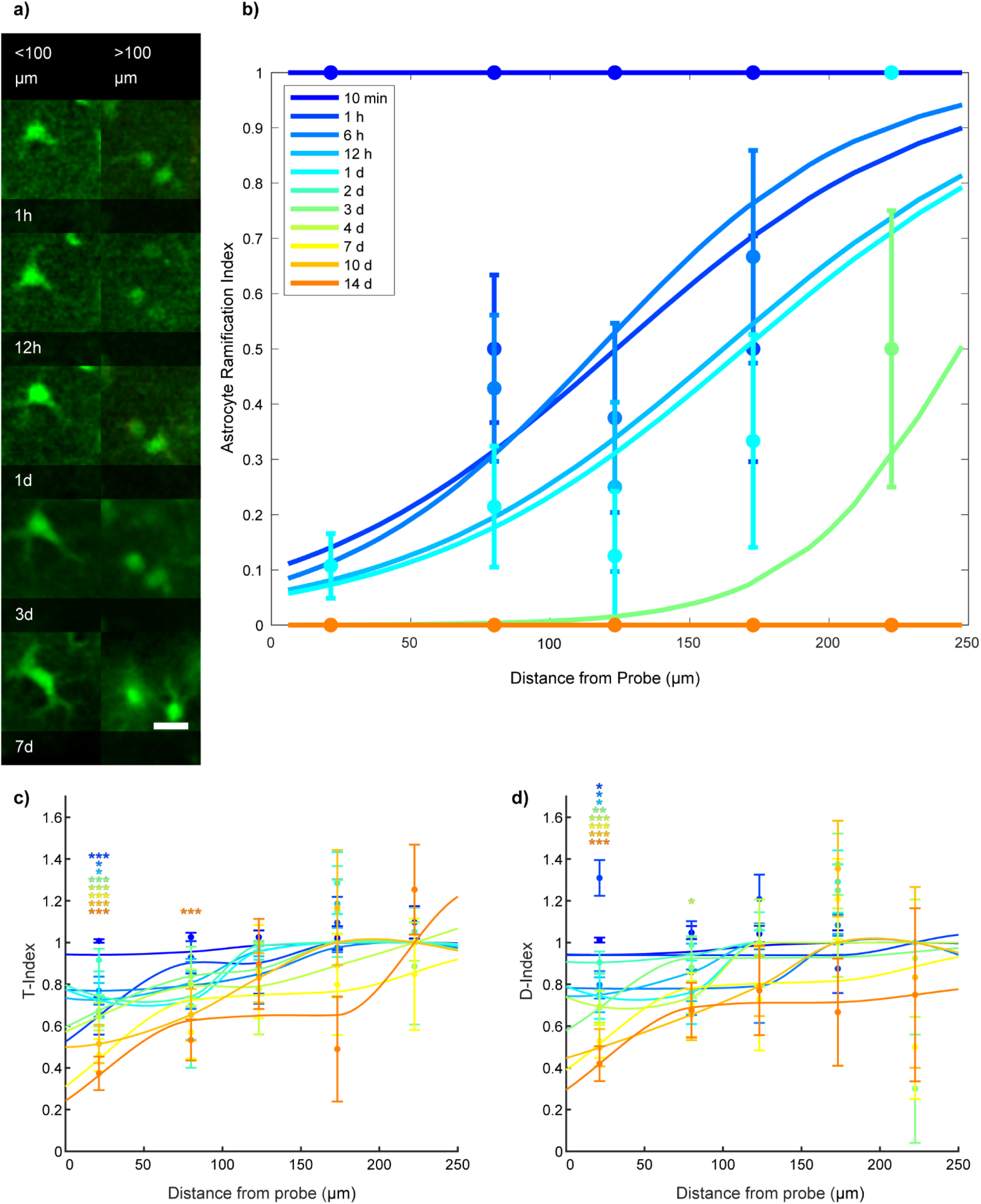
Astrocytes exhibit spatial and temporally distinct patterns of activation following probe implantation. A) Representative images of 20 µm z-projections centered on transitional astrocytes from the probe surface over 7 d post-implantation. Astrocytes within 100 µm from the implant become activated earlier than astrocytes positioned farther from the probe surface (>100 µm). b) Ramification indices of astrocytes as a function of distance from the probe over 14 d post-implantation. By 3 d post-implantation, astrocytes exhibit an activation radius (distance in which there is a 50% probability of ramification) of ∼248 µm from the probe surface. Beyond 3 d post-implantation, virtually all astrocytes demonstrated activated morphology within a 250 µm radius. C) Astrocyte morphological index of its preferred transitional state as a function of distance from the probe (T-index). D) Astrocyte morphological index of its preferred orientation as a function of distance from the probe (D-index). Dunnett’s posthoc test was performed for each distance bin between a control time point (10 min) compared to all other time points. All data shown as mean ± SEM. *p < 0.05, **p < 0.01, ***p < 0.001, color represents time points that are significantly different from the 10 min time point (control).

Given that astrocytes become reactive over distance and time, we compared the morphological changes to the spatiotemporal activation pattern of microglia and NG2 glia. Similar to process migration, astrocytes polarize after microglia, and before NG2 glia. At 2 h and 6 h after probe insertion, astrocytes and microglia demonstrated similar activation profiles (Fig. 6a, b). Microglia were more activated up to 75 µm from the probe surface, while astrocytes were more activated beyond 75 µm. NG2 glia remained ramified during these time points (Fig. 6a, b).12 h post-insertion, astrocytes remained more activated than both microglia and NG2 glia farther than 75 µm from the probe surface (Fig. 6c). At 3 d following insertion, ramified astrocytes could still be observed beyond 200 µm. Astrocytes were still more reactive than NG2 glia at this distance, while NG2 glia themselves were still not completely activated within the observable 250 µm region from the probe surface (Fig. 6d).

**Fig. 6.**
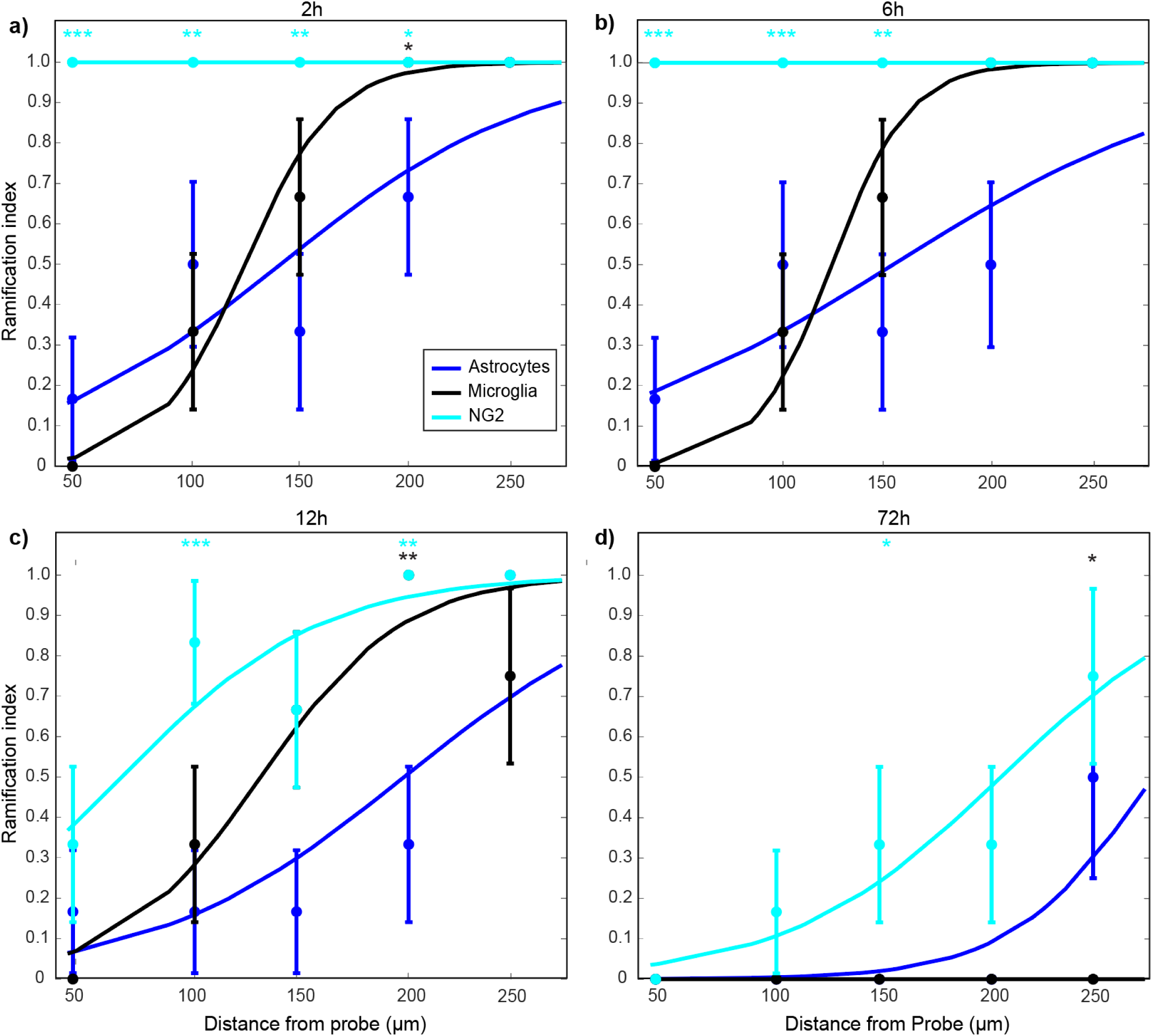
Transitional state patterns of astrocytes differ temporally to NG2 glia and microglia over 3 d following insertion. Ramification of astrocytes, microglia, and NG2 glia are shown as a function of distance from the probe at 2h (a), 6h (b), 12h (c), and 72h (d). By 2 h post-implantation, both microglia and astrocytes are initially activated while NG2 remain ramified. NG2 glia begin to transition by 12 h following implantation by which most microglia and astrocytes are activated up to a 250 µm distance from the probe surface. Information about microglia and NG2 data can be found in the footnote ^2^. * indicates p < 0.05. **p < 0.01, ***p < 0.001, color represents corresponding cell types that are significantly different from astrocytes.

### Astrocyte coverage of the probe begins with soma migration then putative proliferation and differentiation

Although astrocyte process extension, increase in soma area, and morphology could account for the increase in astrocyte area coverage (Fig. 1a), it is possible that astrocyte migration could also contribute to astroglial scarring. The astroglial scarring, which is thought to contribute to device failure [88], could be composed of proliferated astrocytes and NG2 glia differentiated into astrocytes [50, 89], though it is possible that the scar may include some reactive astrocytes that migrated to the injury site. Therefore, we quantified the rate of somatic movement toward the electrode over the first 2 weeks (Fig. 7). Although astrocytes soma movement is stable between 2 – 10 weeks post-insertion [48], astrocyte cell body movements toward the probe began ∼12 h following insertion (Fig. 7a) following process movement (Fig. 2). Astrocyte soma movements were observed at a rate of 0.025 ± 0.025 µm/h, reaching a peak velocity of 0.22 ± 0.023 µm/h on day 3. A significant drop in velocity to 0.08 ± 0.02 µm/h occurs after 4 d and again after 7d (Fig. 7b; *p* < 0.01) remaining stable between day 7 and 14 (data not shown), which may explain why no cell body movements were observed 2 weeks post-implant in previous studies [48]. Similar to the rate of process extension, astrocyte cell bodies exhibited significantly faster movement velocities within 50 μm of the probe surface than further away from the probe (*p* < 0.01). Beyond 150 µm from the probe, astrocyte soma movements were not observed (Fig. 7c), suggesting that astrocytes within 150 µm and particularly within 50 µm may be more likely to contribute to glial scar formation and coordinate a foreign body response. Interestingly, astrocyte movements were significantly far slower than both microglia and NG2, whose movement velocities are at 2.35 ± 0.32 μm/h and 1.1 ± 0.24 μm/h, respectively (Fig 7d; *p* < 0.01).

**Fig. 7.**
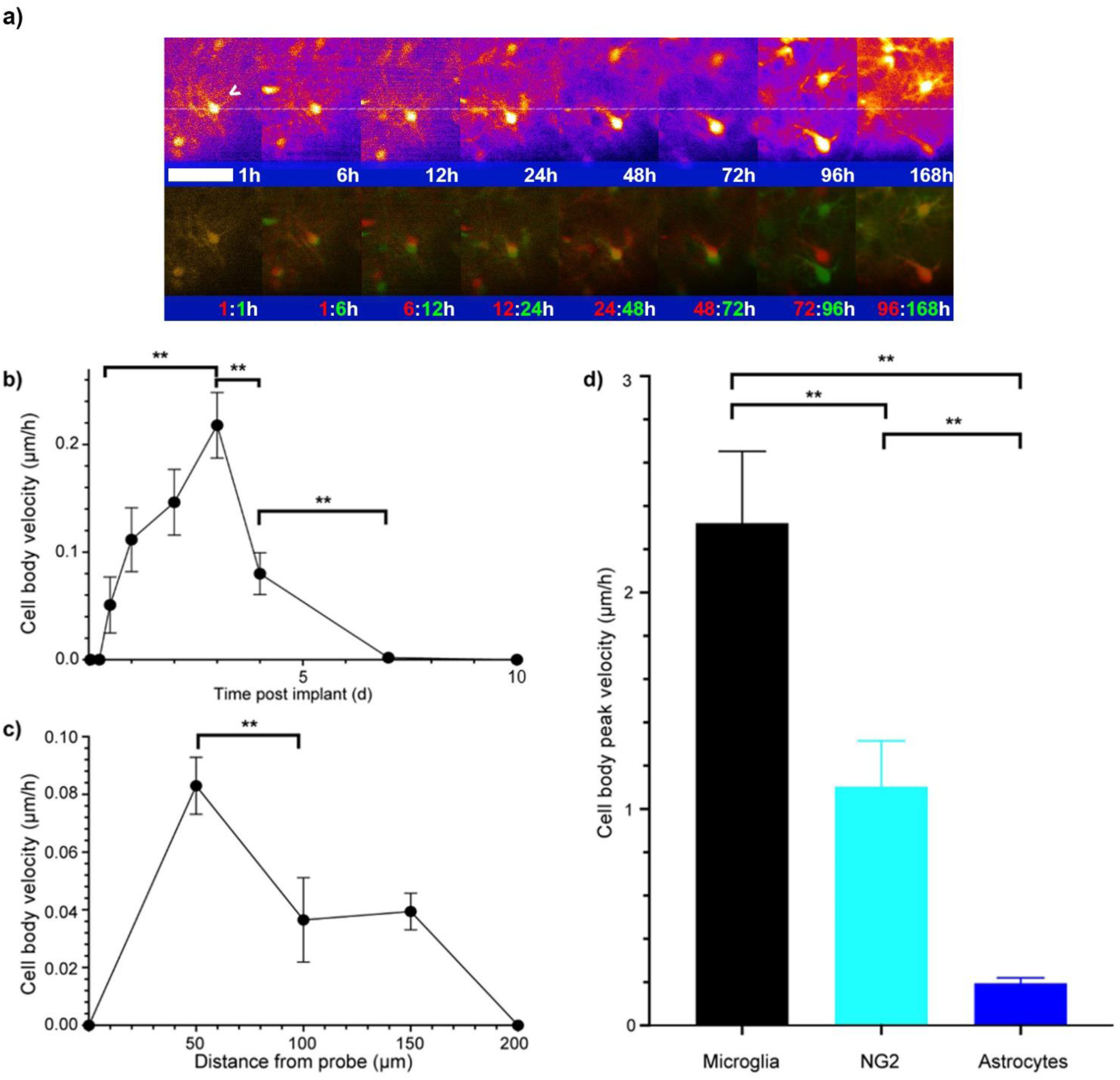
Astrocyte soma movements toward the probe were observed at 12 h post-insertion. A) Superimposed images of 20 µm thick z-projections center on astrocyte soma movement toward the probe surface (*shaded blue*). Red denotes the earlier time point and green denotes the later time point. Yellow depicts overlap between two consecutive time points. Scale bar = 15 µm. b) Astrocyte soma velocities gradually increased and peak at 3 d post-insertion. C) Soma velocity plotted as a function of distance from the probe demonstrate an increased rate of movement for astrocytes more proximal to the probe surface. D) Peak soma movement velocities between glial cells reveal that astrocytes move with the slowest speed compared to microglia and NG2 glia. Information about microglia and NG2 data can be found in the footnote ^3^. * indicates p < 0.05, ** p < 0.01.

Given the limited astrocyte soma movement, which was largely restricted, other mechanisms for increasing astrocyte density around the implant were also examined. Although the astrocyte scarring may include migrated astrocytes, it is believed to be largely formed from newly proliferated astrocytes [90]. Therefore, to identify a potential time point to limit scar formation, we quantified putative astrogenesis (Fig. 8). Putative newly generated astrocytes were characterized as either one of two cellular events: 1) through cell division, observed as the gradual separation of two distinct soma in opposing directions; or 2) through cell differentiation, indicated by a gradual increase in fluorescence intensity within a region of tissue previously lacking GFP+ signal (Fig. 8a). Division and differentiation events were further validated by referencing the 3-D volumetric images in order to rule out false positive classifications due to potential overexposure of fluorescence intensities or drift in imaging plane (Supplementary Fig. 1 & 2).

**Fig 8.**
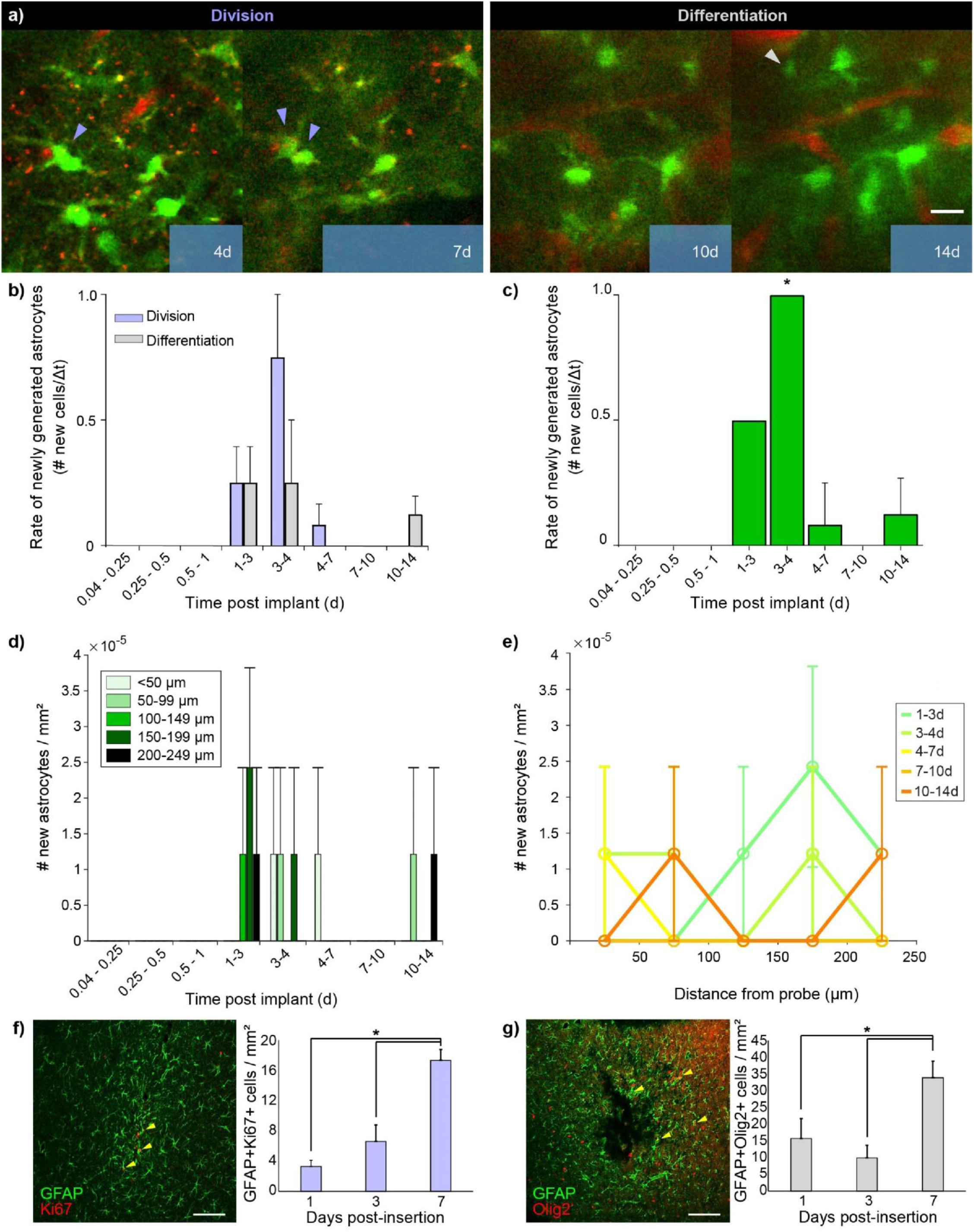
Putative astrogenesis occurs via two distinct cellular processes following probe insertion. ‘Putative’ distinguishes differentiation and proliferation based on *in vivo* GFP data carefully examined in 3D z-stacks over time, in contrast to differentiation and proliferation confirmed through markers of proliferation and differentiation. A) Example 20 µm projection around dividing or differentiating astrocytes. Classification of proliferating astrocytes arising from putative cellular division (left) and/or differentiation (right) near an implanted probe (*shaded blue*). Purple arrowheads demonstrate actively putative dividing cells while the white arrowhead denotes a newly putative differentiated cell. Blood vessels are labeled with SR101 (*red*). Scale bar = 15 µm. b) Rate of putative dividing (purple) versus differentiating (grey) cells per day at different time periods after probe insertion reveal peak in putative astrogenesis at 3-4 d post-insertion. The rate of proliferation was determined by counting the number of new cells in the second time point relative to the first time point that met the classification criteria in A and then dividing this count by the elapsed time. C) Rate of newly generated cells regardless of classification. D) Number of new cells per mm^3^ sorted by distance away from the probe at each period. The new cell’s distance from probe was binned every 50 µm. e) Number of newly generated cells per mm^3^ for different time periods at different distances from the probe. Distances of new cells from the probe were binned at 50 µm. f) Co-labeling for GFAP+ and Ki67+ cells (yellow arrowheads) demonstrates an increase in astrocyte proliferation within first week post-implantation. Scale bar = 100 μm. g) Co-labeling for GFAP+ and Olig2+ cells demonstrates an increase in astrocyte differentiation with first week post-insertion. Scale bar = 100 μm. For all multiphoton analyses, analyzed area = ∼0.1 mm^2^, total depth analyzed per animal = 0.120 mm. Scale bar = 15 µm. All data shown as mean ± SEM, n = 4. \**p*< 0.05

At 1-3 d post-insertion, new astrocytes originated equally from putative dividing and differentiating astrocytes at a rate of 0.25 ± 0.144 new cells/day over the field of view (Fig. 8b). At 3-4 d post-insertion, dividing astrocytes appeared a rate of 0.750 ± 0.250 new cells/day, while putative differentiating cells occurred a rate of 0.250 ± 0.250 new cells/day, showing a higher trend in putative astrogenesis from dividing cells relative to putative differentiating cells (*p* = 0.625). Astrocyte generation slowed down after 4 d with no significant differences between putative division and differentiation (Fig. 8b). When observing the combined rate of putative astrogenesis from both putative dividing and differentiating astrocyte populations, the rate at 3-4 d was significantly higher than the rate between 0.04-0.25 d (within first 6 h) (Fig. 8c, *p* < 0.05). Distribution of new astrocytes at 1-3 d post-implantation revealed putative astrogenesis occurred primarily beyond 200 µm from the probe surface (Fig. 8d). However, at 3-4 d a decrease in the number of newly formed astrocytes was observed beyond 100 µm from the probe, while an increase in putative astrogenesis occurred within the first 100 µm. Between 4-7 d, putative astrogenesis was observed only within the first 50 µm and by 7-10 d no new astrocytes were observed within 250 µm from the probe. However, new astrocytes were found at 50-99 µm and 200-249 µm from the probe surface during 10-14 d (Fig. 8d and 8e).

It should be noted that the results exhibit a large range of variability due to a low rate of observed putative astrogenesis. A repeated-measures two-way Friedman test indicated a difference among one of the groups when comparing time points (*p* = 0.0442), however, no statistically significant differences were observed in the number of astrocytes per mm^3^ between each time period when compared to the control period (0.04-0.25 d) using Dunn’s multiple comparison test. To further validate the generation of new astrocytes around an implanted microelectrode, we performed additional immunohistochemical analyses to visualize and quantify the number of proliferating and differentiating astrocytes. Co-labeling for GFAP+ and Ki67+, a marker for cellular proliferation [91, 92], revealed a significant increase in the number of proliferating astrocytes up to 7 d post-insertion (Fig. 8f, *p* < 0.05). Co-labeling for GFAP+ and Olig2+, a marker for astrocyte differentiation, demonstrated a significant increase in differentiated astrocyte cells similarly within the first week post-insertion (Fig. 8g, *p* < 0.05). These results are in agreement with previous post-mortem immunohistochemical studies [60].

So far, we demonstrated that after electrode insertion, astrocytes extend processes and begin polarizing and migrating toward the electrode, become hypertrophic, and potentially begin proliferating and differentiating. However, it is the coverage of the electrode that has been suggested to influence recording performance. Therefore, we quantified the area of the electrode that was covered by dense astrocytes (Fig. 9). The percentage of astrocytes covering the surface of the probe significantly increased from 14 ± 1.2% to 17 ± 2.3% (Fig. 9b; *p* < 0.05). This change in surface coverage was markedly lower in comparison to microglia and NG2 glia surface coverage from 52.8 ± 4.5% to 91.4 ± 3.3% and 36.3 ± 4.2% to 72.6 ± 8.9%, respectively [50]. Probe coverage by microglia and NG2 glia was previously only quantified up to 72 h post insertion, therefore we compared the astrocyte probe coverage at that time. Additionally, a significant increase in astrocyte density over time can be observed in the region of tissue adjacent to the probe (Fig. 9c, d; *p* < 0.05). While a nominal increase was observed within 100 µm of the probe surface as early as 24 h, a larger number of astrocytes appeared across the entire 250 µm imaging range by 3 d following implantation. In contrast to 4 d, astrocyte density at 7-10 d at distances beyond 200 µm were lower yet there was an increase in density at distances less than 100 µm from the probe. By 14 d astrocyte density was evenly distributed in the first 200 µm from the probe (Fig. 9d).

**Fig. 9.**
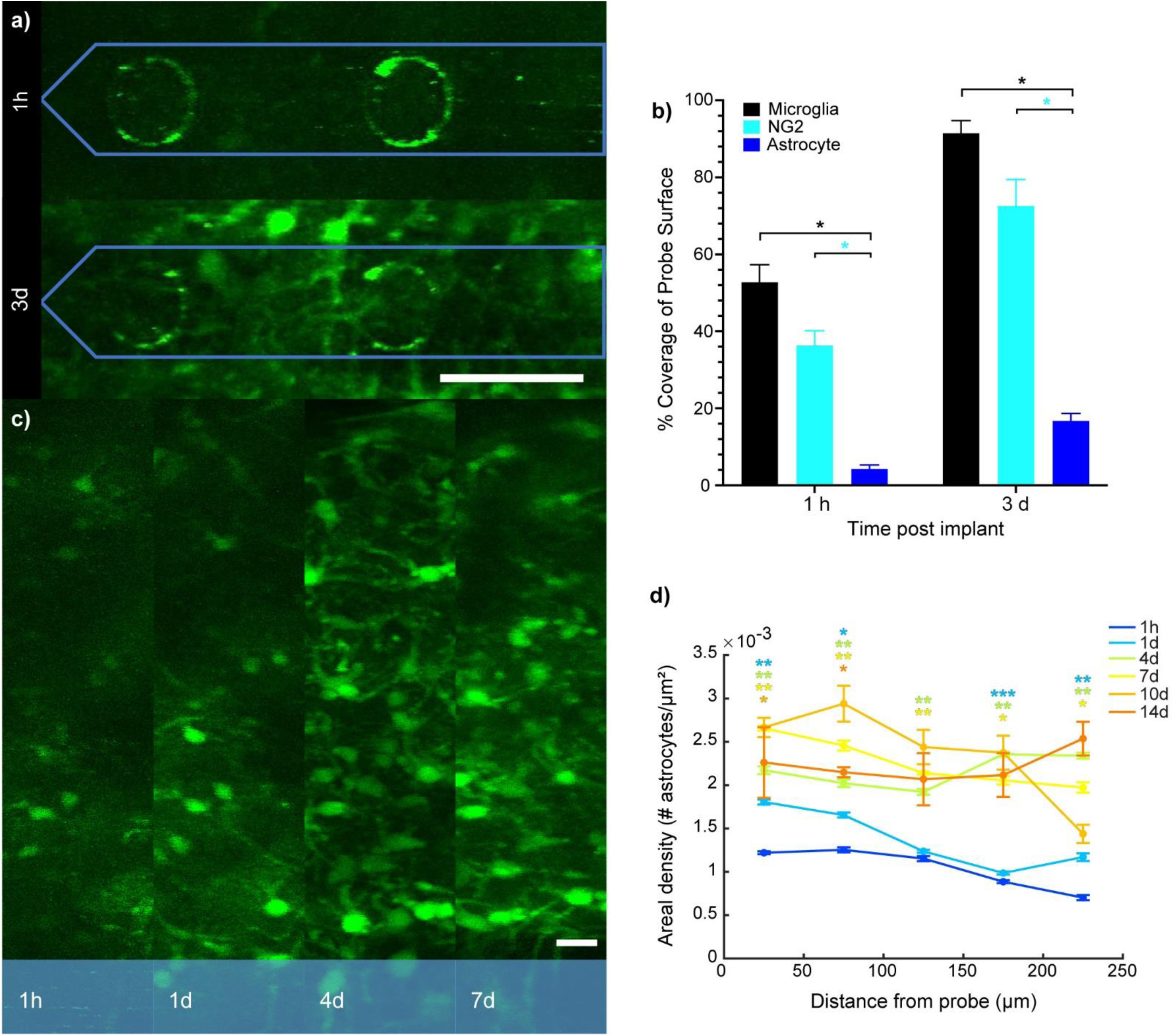
Increased astrogliosis and surface encapsulation of implanted probes. A) GFP+ signal intensity within astrocyte processes encapsulating the probe (*blue outline*) increased from 2 h to 3 d post-insertion. Scale bar = 50 µm. b) Percent of coverage within 20 µm of the probe surface at 1 h and 4 d by different glial cells demonstrate minor contribution by astrocytes. C) Increases in astrocyte distribution adjacent to probe surface (*shaded blue*) at 1 h, 1 d, 4 d, and 7 d post-insertion. Scale bar = 15 µm. d) Astrocyte density of astrocytes surrounding the probe as a function of distance from the probe at 1 h, 1 d, 4 d, 7 d, 10 d and 14 d post-insertion. Information about microglia and NG2 data can be found in the footnote ^4^. * indicates p < 0.05, ** p < 0.01, *** p < 0.001, color represents time points that are significantly different from the 1 h time point (control).

Together, the astrocyte process extension and migration toward the electrode, as well as the somatic hypertrophy suggest that astrocytes begin their reactive response to electrode implantation injury within 30 minutes and continue to respond though process polarization, putative proliferation and probe coverage up to 14 d after implantation. Importantly, although astrocyte migration and process extension are slower than microglia and NG2 glia, the astrocyte response begins after microglia and before NG2 glia suggesting there could be a coordination between these cell types.

## Discussion

BCIs enable both the decoding of neural signals from the brain [93] and stimulation or neuromodulation for clinical therapy [94]. However, depreciations in recorded signal over time increase the difficulty of extracting useful spiking information from the noise floor, which impedes the widespread use of this technology. Device implantation ruptures surrounding brain tissue and mechanically strains nearby neurons, preventing them from firing properly [28, 31, 45, 46]. Besides direct injury due to insertion, neuroinflammation and foreign body responses from chronic device implantation result in astroglial scarring around the neural implant and neurodegeneration (see review [95]), which ultimately impact long-term recording and stimulating performance [5, 28, 31]. Astrocyte activation and glial scar formation is a well-established and robust feature of the biological response to intracortical implants [96]. However, a previous study using two-photon microscopy demonstrated that the expression of ALDH1L1-GFP in astrocytes remain largely the same from 2 to 10 weeks post insertion [48]. Therefore, it remained unclear when astrocytes form the astroglial scar around implanted arrays, suggesting that a substantial proportion of astrocyte reactivity and scar formation dynamics may occur within the first 2 weeks. To fill in the gaps on the dynamic astrocytic activity within this early implantation period, we imaged transgenic mice (ALDH1L1-GFP and GFAP-GFP) longitudinally using two-photon microscopy to observe spatiotemporal changes in astrocyte morphology and behavior around an implanted device. Aldh1l1 promotor allows to reliably label astrocyte cell bodies and processes in cortex, while GFAP, in contrast, primarily labels astrocytes with low expression levels until prominent neuroinflammation occurs. Therefore, we use Aldh1l1 promotor to clearly observe astrocyte dynamic response during acute insertion, and switch to GFAP promotor to visualize astrocyte involvement during later insertion-induced neuroinflammation when the upregulation of GFAP strengthens the visualization of astrocytes. Previous immunohistochemical data have confirmed that astrocyte labeling GFAP are all positive for Aldh1l1 in cortex [97], which indicates that our observation of GFAP+ activity at later implantation stage could be consistent to acute Aldh1l1 response. Our study details the spatiotemporal dynamics of astrocyte reactivity and putative proliferation as well as their behavior in relation to the activity of local blood vessels during the first two-week device post-implantation. Additionally, we highlight temporal comparisons between astrocyte reactivity within that of microglia, NG2 glia, and vascular dilation. We demonstrate that astrocyte activity is highly dynamic during the acute-to-chronic period around implant probes and provide new insights into the cascade of biological events at the device-tissue interface.

### Astroglial scarring and its contribution to impedance of neural recording

The biological cascades initiated by probe insertion can play a major role in disrupting the electrical properties of the neural interface [24, 28, 31, 95, 98, 99]. For example, the resulting microglial scarring and probe encapsulation forms a physical diffusion barrier that impedes electrical signal transmission between neurons and microelectrode contact sites [53, 95, 99–101]. Similarly, astrocytes can also contribute to the encapsulating glial scar via expansion of cellular membranes [34, 102] and thus potentially increase impedance of the device. Injury can cause hypertrophy of reactive astrocytes, increase in their volume, and dramatic upregulation in GFAP expression over the implant region [57, 83]. While it is known that these morphological changes occur by 3 d after injury [81, 103], the early spatiotemporal dynamic morphological changes of astrocytes around implanted devices remained uncharacterized. A more comprehensive understanding of the temporal dynamics of astrocyte reactivity following electrode insertion in the context of other cellular responses could help to identify critical time periods for therapeutic intervention and improve long-term device performance.

Intracortical microelectrodes induce neurovascular [47] and axonal damage [72] immediately upon insertion. Release of tissue injury signals from locally ruptured blood vessels or damaged neurons are known to activate surrounding microglia and astrocytes following brain injury (see review [95]). Astrocytes become activated in response to cytokines and chemokines secreted from surrounding glia, including activated microglia (see review [104]). In this work, we demonstrated that astrocytes extend their processes as early as 1 h post-implant (Fig. 2b), which trailed microglia process extension but not NG2 glia (Fig. 2d). This supports the idea that the initial stages of astrocyte reactivity occurs in response to cytokines released by microglia [105]. Importantly, reactive astrocytes bidirectionally communicate with microglia and can release molecules such as MCP-1 [106], CCL2 [107], IP-10 [108] and CXCL10 [109, 110] to modulate microglial reactivity and mobility [111, 112]. For example, MCP-1 has been shown to be especially synthesized in hypertrophic astrocytes [106] such as the hypertrophic astrocytes we observed (Fig. 3). This MCP-1 can induce microglial proliferation [113] as well as migration of amoeboid microglia [114]. Microglia can also form tight junctions with each other [101], which act as a diffusion barrier and inhibit the transmission of neural signals, leading to an increase in electrochemical impedance [95, 102]. Since astrocytes begin to show signs of reactivity within an hour of implantation but peaked between day 2-4, blocking astrocyte production or release of MCP-1 during this time frame may be beneficial to reduce neuroinflammation and promote long-term device performance. While the communication between astrocyte and microglia with NG2 glia are less understood, the sequential order of glial activation and migration suggest a coordination among different glial cells in the foreign body response.

Following process extension toward the implant, reactive astrocytes became hypertrophic (Fig. 3; peak day 2-4) and were shown to have movements toward the injury site (Fig. 7b, c; peak day 3). It is possible that astrocytes become hypertrophic and migrate toward the probe in an attempt to mitigate vascular leakage by producing a diffusion barrier via tight junction formation [102]. This in turn would make it less permissive for cells and plasma proteins to leak through the BBB [115], which negatively impacts impedance. In addition to astrocyte movement toward the implant (Fig. 7), putative astrogenesis was also observed around the injury site (Fig. 8). Astrocytes can proliferate through self-division and from differentiation of local progenitor cells, such as NG2 glia [116], however, this only occurs when there is significant damage. Consistent with this, some astrocytes appeared to be dividing around the injury region. These putative proliferating astrocytes maintained high GFP+ signal throughout the division process (Fig. 8a). Additionally, some astrocytes spontaneously appeared around the implant and did not divide from any observable GFP+ cells nearby (Fig. 8). The fluorescence intensities of these astrocytes started out noticeably dim before increasing in intensity over days suggesting differentiation of precursor cells into astrocytes, potentially from nearby un-labeled NG2 glia that migrated closer to the implant surface during this time [117, 118]. Origin of NG2 cell-derived reactive astrocytes are dependent on the type of CNS injury and therefore may serve different pathological functions compared to normally dividing astrocytes [119]. Immunohistochemistry in animals sacrificed 3 days and 7 days post-implantation further confirmed the low levels of astrocyte division (GFAP+/ki67+) and differentiation (GFAP+/Olig2+) around chronically implanted microelectrodes (Fig. 8f-g). Interestingly, putative astrogenesis primarily occurs >200 μm from the implant, but this shifts to <100 μm by day 3-4 post-insertion (Fig. 8c) and is temporally correlated with the peak of movement velocity (Fig. 7b), suggesting newly generated astrocytes may migrate toward the implant with a high velocity. This increase in astrocytes near the implant may indicate the beginnings of glial scar formation [120].

It should be noted that the measurements of astrocyte soma movements may capture more than soma migration (Fig. 7). As we observed, peak soma movement velocity (Fig. 7b) coincides with the largest soma area (Fig. 3b) and the largest vessel diameter dilation (Fig. 4b). Both tissue swelling and the blood vessel dilation may result in the displacement of surrounding tissues. Our current method cannot exclude the possibility that the displacement of surrounding tissues also contributes to the observed movements of astrocyte soma. Notably, the movement of astrocytes around the implant were far slower than microglia or NG2 glia (Fig. 7d). Nevertheless, multiple proteins could contribute to astrocyte migration, including Cx43 [121], Cdc42 [122] and DCLK [123]. Future studies to knockout these proteins could help to understand the mechanisms underlying the astrocyte soma movements after the device insertion and identify therapeutic strategies to improve device integration with the brain.

Surface coverage within 20 µm of the electrode was less for astrocytes than microglia (Fig. 9b, *p*<0.05), consistent with previous reports [60, 124]. This is supported by the fact that microglia activation is greater (ramification index is closer to 0) than astrocytes and NG2 glia within 50 μm of the implant (Fig. 6a, b) and both astrocyte process and soma velocity is slower than microglia and NG2 glia (Fig. 2d, 7d). However, astrocyte encapsulation is more pervasive at further distances away from the probe (Fig. 2c), which may potentially contribute more to the outer layer of reactive scar tissue. These results illustrate coordination between microglia, NG2 glia, and astrocytes in forming the inner (microglia & NG2 glia) and outer (astrocyte) layer of reactive scar tissue around implanted probes and are confirmed by post-mortem immunohistochemical studies [60].

Glial scarring has long been considered to affect the neural recording performance [5, 88, 99, 125]. Efforts have been made to reduce astrocytic encapsulation and avoid the increase in the impedance of neural recording [126–128]. However, a modeling study suggests that eliminating glial scarring may not be effective in improving device-tissue interface [129], although this study did not exclude material failure in the data used to generate the model [23, 130, 131]. It should be noted that this astrocyte reactivity is not solely about glial scar formation, but astrocyte activity also crucially impacts axonal regeneration [132], reduces the mechanical strain [133, 134] as well as prevents the spread of inflammation (see review [57]). Therefore, it is important to first understand the time course of astrocyte reactivity to device implantation in order to begin identifying therapeutic targets that can balance tissue health with an increase in impedance for neural recording.

### Astrocyte-mediated alterations in single-unit amplitude

Implantation-induced astrocyte reactivity may also affect recorded single-unit amplitudes. Single-units represent the activity of individual neurons extracted from extracellular recorded potentials used in neuroscience research [135] and neural prosthetic control [93], and may provide information in a clinical setting [136]. Since single-unit waveforms are combined with background electrical noise, a higher recorded single-unit amplitude is more conducive to the separation of single-unit waveforms from that noise. Importantly, single-unit amplitudes decrease after device insertion [5, 28, 137], partly due to the increase in the impedance caused by scar tissue (as described above). However, there may be other biological factors that may contribute to the reduced single-unit amplitudes.

For example, astrocyte reactivity coincides with an increase in the release of cytokines that leads to progressive neuroinflammation and ultimately neurodegeneration [138, 139]. We demonstrate that reactivity signatures such as process elongation begins within 1h following device implantation (Fig. 2b) and drops shortly after. This may suggest a transition from debris engulfment [140, 141] toward neuroinflammation. In support of this, a recent transcriptional study reported an increase in the expression of complement factor 3 (C3) within 4 h after probe implantation [142]. Since astrocytes can release C3 via the NF*κ*B pathway [143], it is possible that activation of astrocytes between 1 h and 4 h (Fig. 2b and Fig. 5b) may play a role in the release of C3, which may contribute to neuroinflammation [144–146]. Importantly, neuroinflammation around the electrode site may prevent neurons from firing properly or cause them to undergo apoptosis [28, 31, 45, 46]. Thus, if the astrocyte reactivity transitions to a stage to promote neuroinflammation and neurodegeneration near the electrode, there would be fewer neurons within the optimal recording distance and the action potential waveform would have to travel farther through the tissue resulting in attenuated signals.

Alternatively, astrocyte reactivity may also cause pressure within the extracellular space, which reduces single-unit amplitudes. Both hypertrophy (Fig. 3) and proliferation (Fig. 8) indicate astrocyte reactivity and an increase in extracellular pressure [147]. Increased pressure within the tissue may lead to the deformation of axons, resulting in abnormal sodium influx [148]. In addition, mechanical strain has been shown to reduce the peak inwards of sodium currents [149]. These studies suggest mechanical strain will lead to less sodium influx during action potentials, potentially reducing single-unit amplitudes. Therefore, the mechanical strain places on neuron somata and axons due to hypertrophic astrocytes and proliferating glia could impair proper ion channel dynamics and reduce single unit amplitude.

Finally, astrocyte reactivity may exacerbate the mismatch of metabolic supply and demand. ATP in the axons is essential for maintaining the ionic gradients and firing of action potentials [150]. During device insertion, axons are inevitably affected either directly by injury [31] or deformation (as discussed above). Progressive inflammation may cause mitochondrial damage and decreased ATP production [151]. On the other hand, repair of damaged tissue and phagocytosis of dead cells increase the ATP demand [152, 153]. The diameter of blood vessels increases significantly after insertion (Fig. 4, *p*<0.05), suggesting a need to satisfy energetic demands with increases in blood flow. Normal astrocytes can store glycogen to supply energy for neuronal firing [154]. However, the progression to reactive astrocytes after insertion (Fig. 5) may disrupt the metabolic supply chain via astrocyte-neuron [155] and astrocyte-oligodendrocyte-neuron [156] connections. If this connection is disrupted, the energy supply to these cells and the level of available ATP in the axons could be reduced. Importantly, reduced ATP in optic nerves due to high intracranial pressure reduce the amplitude of the compound action potential [157]. Therefore, the single unit amplitude could also be reduced if astrocyte reactivity does in fact lead to the impairment of energy supply to neurons.

### Astrocyte-mediated alterations in neuronal firing activity

Detecting changes in neuronal activity is essential for decoding brain activity in BCI applications as well as neuroscience research. Recent literature suggests that astrocytes actively modulate neuronal firing patterns [158, 159]. However, alteration of astrocyte function following injury may have a profound impact on the regulation of local neural circuits [160] and neurovascular coupling (see review [161]). Therefore, it is necessary to understand the changes in astrocytic function during the course of implantation and their impacts on local circuit activity in order to develop targeted intervention strategies that guide the development of future clinical therapies.

Previous electrophysiological studies observed increase in neuronal firing within the first 24 h post-insertion [5, 29, 137] suggestive of hyperexcitability within nearby neurons. The combined mechanical forces induced by astrocyte hypertrophy (Fig. 3) and an accommodation of the probe volume by the cortex may effectively compress the extracellular space, increasing the concentration of neurotransmitters and ions within the local tissue microenvironment [162]. For example, the increased concentration of glutamate [163, 164] and potassium [165] have the potential to induce the hyperexcitability of surrounding neurons. However, it is important to note that we cannot exclude the possibility that increased firing rate comes from recovery following anesthesia over the course of the first 24 h after device implantation surgery [9].

Neuronal activity then decreases from 24 h to 48 h post-implantation [5, 29, 137], which indicates a potential reduction in excitatory synaptic transmission. Astrocytes are known to increase the excitability of excitatory synapses by releasing of glutamate or decreasing glutamate uptake [166–170]. However, after implantation, astrocytes extend their processes (Fig. 2), encapsulate the probe surface (Fig. 9), and enter a reactive and hypertrophic state (Fig. 3, 5). This suggests that astrocytes would have fewer processes available to promote excitability of local excitatory synapses, leading to a reduction in neuronal firing. On the other hand, astrocytes can also increase the activation of local inhibitory interneurons [171]. In this case, loss of inhibition due to lack of astrocyte activation of inhibitory synapses may lead to increases in excitatory activity, thereby increasing neuronal signaling. However, this possibility seems to contradict the decreases in firing rate observed during the acute period following insertion [5, 29, 137]. Nevertheless, ongoing inflammation and neurodegeneration may exacerbate the decreases in neuronal firing rates and its impact may be highly time course dependent. For example, the astrocyte activation (Fig. 5) coincides with increased inflammation, upregulation of pro-inflammatory cytokines, and changes to osmotic pressures, which can also lead to silencing of local neural activity [31].

In previous studies, visually-evoked multi-unit firing rate could increase again during the first week starting from day 3 post-insertion [5, 29, 137]. It is possible that the effect of increased concentration of glutamate and potassium overcome other factors since the soma area of astrocytes increase dramatically and significantly during this time (Fig. 3, *p*<0.05). In addition, proliferation rates within astrocyte are significantly higher during day 3 to day 4 (Fig. 8c, *p*<0.05), suggesting more astrocyte processes are available to promote excitability of local excitatory synapses, as discussed previously [166–168]. This rising trend in neuronal firing rates does not last long, however, and maintains a downward trajectory beyond 4 d post-insertion [5]. Yet, astrocyte-mediated process extension (Fig. 2), soma movements (Fig. 7) and hypertrophy (Fig. 3) appear to decrease during this period suggesting that future research should explore other factors that may contribute more to the decline in recordable neuronal activity, such as degradation of myelin sheath for signal conduction [72], dysfunction within the blood-brain barrier, and/or neurodegeneration.

While astrocyte signaling modulates local neuronal firing activity [172], coordinated activity of these functionally altered neurons and synaptic circuits possibly affect even wider range of information processing in high-order brain function. Recently astrocytes have been recognized as a critical mediator of neuronal oscillation across large-scale brain areas [173–175], as specific knockout of astrocytic GABA_B_ transmission leads to power reduction in gamma band [175]. Previous electrophysiological recording in mice reveals severe fluctuation in gamma relative power in the first two weeks after implantation [5], which may temporally correspond to our observation in active astrocytic process activity (Fig. 2), soma movements (Fig. 7), gradual encapsulation (Fig. 9), and population turnover (Fig. 8). Astrocytes can target gliotransmitter release onto many dendritic and axons that are potential for synchronization of neuron ensembles [176–178]. Therefore, the observation of hypertrophic state and convergence of remaining processes toward the probe (Fig. 5) following implantation suggests less available processes in proximity to dendrites and axons to achieve selectively release, resulting disordered astrocytic modulation on network coordination in high-frequency range. Future studies should investigate whether the activation of astrocytes will cause the highly fluctuated, generally decreasing power in gamma band over two weeks insertion [5]. Meanwhile, our result shows the activated state of astrocytes is up to 250 μm away from the probe at day 14 post insertion. Together, these findings motivate future investigation to examine the effect of microelectrode implantation on disturbance of astrocytic signaling on large-scale network oscillatory rhythms that carry information transmitted to a distinct functional brain region.

### Astrocytes support of neuronal activity indirectly via coupling with vasculature and oligodendrocytes

Astrocytes are an integral component of the neurovascular unit and are considered indirect regulators of neuronal metabolic activity by facilitating the bidirectional exchange of modulatory signals between neurons and blood vessels during neurovascular coupling [179]. Previously, we have demonstrated that local blood vessels expand within the first 3 days following device implantation [50]. Here, we show that these changes in diameter are positively correlated with changes in soma area of adjacent astrocytes near implanted probes (Fig. 4c). Astrocytic endfeet completely encapsulate the entire abluminal surface of blood vessels within the brain, allowing astrocytes to regulate important homeostatic processes such as cerebral blood flow, blood-brain barrier permeability, and the exchange of nutrients and solutes across the endothelium [180, 181]. It is possible that astrocytes compensate for an increase in endothelial surface area by expanding their cellular membranes to continue supporting these critical regulatory mechanisms. Alternatively, astrocytes are known to play an active role in repairing blood vessels following vascular injury. For example, selective ablation of astrocytes results in disrupted vascular repair following stroke [182]. Hypertrophy as a form of astrogliosis could be indicative of a neuroprotective response to direct vessel rupture or increased vascular permeability, both of which are known to be direct consequences of immediate and prolonged device implantation [47, 183]. We also observed that astrocyte proliferation peaks during this window of maximal astrocyte and vascular reactivity (Fig. 8c). Astrocytes are highly proliferative within juxtavascular sites after acute injury [184]. Depending on the severity of vascular injury, an over proliferation of astrocytes may be an indication of reactive scar formation (see review [57]). It is still unclear whether scar formation is beneficial [132] or detrimental [185] to the functional outcomes of perilesional brain tissue and the consequences of astrocyte activation on both vascular dynamics and neuronal function around chronically implanted devices remain to be seen.

Furthermore, the co-occurrence of astrocytic hypertrophy and vessel dilation may be due to alteration of molecules and ions in the extracellular space, such as nitric oxide [186, 187] and potassium [164, 188], which is potentially related to blood-brain barrier dysfunction and neurodegeneration after insertion [60]. Astrocytic endfeet predominantly express aquaporin 4 (AQP4) channels, which facilitate the transport of water into and out of the brain, as well as inward rectifying K^+^ channels (Kir4.1), which are responsible for regulating potassium ion concentrations within the extracellular space [189]. During seizure conditions of reactive gliosis and neural hyperexcitability such as in epileptic brains, AQP4 channels redistribute from perivascular to perisynaptic compartments within astrocyte processes and consequently result in impaired K^+^ clearance from the extracellular space, increasing neuronal sensitivity to concurrent seizures [164, 190]. While it is unclear from the results of our study whether the astrogliosis observed around blood vessels adjacent to the implant suggest impaired potassium ion regulation, it could provide insight into a causal relationship between vascular responses, astrogliosis, and the disruption of firing activity within local neural circuits surrounding chronically implanted microelectrode devices and is worth further study.

Besides coupling with vasculature, astrocyte also modulate neuronal functionality through coupling with oligodendrocytes. While neurons cannot store glucose as glycogen, astrocytes uptake glucose from the vessel, digest it into small-molecule intermediate, and shuttle these metabolites toward neurons through oligodendrocytes that have large contact area with axons [156]. Therefore, astrocyte-oligodendrocyte coupling is mediated, in part by gap-junctions, and is a vital mechanism to meet neuronal energy demand [156]. However, during chronic implantation, the dynamic astrocytes process (Fig. 2) and soma movements (Fig. 7) are likely to dissociate from immobile oligodendrocytes, damaging the metabolic support to axons (and oligodendrocytes) that need substantial energy to maintain normal activity. This may explain the early loss and silencing of neuron firing activity [5, 31]. In addition, oligodendrocytes are vulnerable to neuroinflammatory environment [60, 72] and may require energy to resist degeneration from injury and maintain normal functionality. The observation that astrocyte division mainly occurred at 24 h to 7 d suggests energy utilization prioritizes themselves rather than supporting oligodendrocytes or neurons, which matches the early start of myelin loss [72] and neuron apoptosis [60] reported in previous publications. Moreover, the aggregation of reactive astrocytes near the probe and limited proliferation from 7 d to 14 d suggest that fewer ramified astrocytes are available for rebuilding astrocyte-oligodendrocyte metabolic coupling, which could coincide with severe demyelination and in turn, loss of neuronal viability during chronic implantation [60, 72]. Together, these findings motivate future studies further investigating metabolic mechanisms that link these temporally correlated cellular changes following device implantation.

### Biomaterial-based approaches to modulate astrocyte behavior

Stiffer substrates, such as with traditional silicon electrodes (∼10^2^ GPa) used in this study, elicit a much larger mechanical mismatch when inserted into softer brain tissue (∼10^-5^ GPa) and are more likely to exacerbate neuroinflammation compared to more mechanically-compliant materials due to repeated insults to the brain arising from micromotion [191, 192]. Astrocytes especially are highly sensitive to changes in mechanical cues within their local microenvironment [193]. For example, GFAP+ immunoreactivity within astrocytes is significantly attenuated in the presence of more compliant devices compared to stiffer probes [194]. Biomaterial-based strategies to improve the softness and/or flexibility of intracortical devices have been explored to modulate glial activation and scar formation, such as use of more elastic materials including conductive polymer coatings or hydrogels [24, 195]. However, these strategies have their own shortcomings. For example, flexible devices are more difficult to insert into the brain and often require the assistance of an insertion shuttle [52, 196] or use of more innovative, mechanically-adaptive materials that respond dynamically to the tissue microenvironment following implantation [197–199]. Furthermore, an alternative and/or combinatorial strategy to modulate astrocyte responses are fabrication of devices with smaller feature sizes with the goal of minimizing glial activation by reducing the amount of contact area between glial cells and the device surface [71, 200]. Probes with varying geometries, such as multi-shank electrodes of varying dimensions as well as arrays fabricated with lattice-like architectures, demonstrate differences in microglia and astrocyte activation very clearly [52, 201]. Finally, untethered, or ‘floating’, electrode arrays have demonstrated favorable tissue response outcomes over probes tethered to the skull due to diminished micromotion-induced neuroinflammation and overall smaller device footprint, which can favor long-term device fidelity and performance [70, 202].

Electrode surface coatings are also emerging as promising biomaterial interventions that can modulate glial inflammation and promote neuroprotection via the controlled release of therapeutic drugs or mobilization of bioactive substances to the implant surface. Promising pharmacological candidates include dexamethasone, an anti-inflammatory drug which suppresses scar formation by modulating cellular process dynamics following device insertion *in vivo* [76], and minocycline, an antibiotic which can reduce microglia and astrocyte activation, enhance neuronal survival, and improve device recordability [203]. Resveratrol, an anti-oxidative drug, can preserve neuronal densities while attenuating astrogliosis around chronically implanted microelectrodes by preventing accumulation of harmful reactive oxidative species (ROS) within the tissue microenvironment [203]. More recent advances in understanding the intrinsic mechanisms governing astrocyte activation and proliferation have identified potential therapeutic targets of interest. For example, selective ablation of Na+/H+ exchanger isoform 1 (NHE1) in astrocytes leads to improved neurological outcomes by reducing astrogliosis, preventing vascular damage, and enhancing neuroprotection following ischemia [204]. Cariporide (HOE-642) is a drug that can inhibit NHE1 activity and has recently shown to modulate glial activation and surface encapsulation following device implantation *in vivo* [205]. Modification of electrode surfaces with dissolvable hydrogels or drug-eluting polymer coatings can allow each of these drugs targeting glial activity to be experimentally secreted in a controlled manner. Finally, the use of electrically-conductive polymer coatings such as poly (3,4-ethylenedioxythiophene) (PEDOT) demonstrate multiple biological advantages for improving electrode-tissue integration such as 1) providing a more mechanically-favorable substrate for glial attachment, 2) improving neural recordability through enhanced material conductivity, and 3) facilitating controlled release of therapeutic drugs encapsulated within electrostatically-adhered nanoparticles via electrical stimulation [206–208].

Furthermore, the brain’s capacity for neurogenesis is limited. Therapeutic approaches aimed at enhancing neuroregeneration after brain injury have garnered increasing interest, particularly following the discovery that previously non-neurogenic glial cells in the intact brain can display neural stem cell-like features under certain pathological conditions [209]. For example, astrocytes demonstrate multipotency and self-renewing behavior *in vitro* when isolated from injured CNS tissue suffering from traumatic brain injury as well as the capacity to form neurospheres following cortical stab wound injury or cerebral ischemia [210]. This phenomenon is injury-dependent since similar astrocyte plasticity is not observed within an Alzheimer’s model of chronic amyloidosis [210]. The therapeutic implications of these discoveries have led to investigations aimed at directly reprogramming reactive glial cells *in vivo* following brain injury via pharmacological/genetic manipulation or exposure to different transcription factors [211–213]. It is possible, using the biomaterial approaches described previously, that these reprogramming factors can be experimentally released from the site of device implantation into the local tissue microenvironment in both a predetermined and controlled manner. By observing the timeline of astrocyte proliferation and movements noted in this study to strategically inform of a therapeutic window of drug delivery (i.e. around day 3-4 post-insertion), one could theoretically optimize the reprogramming of activated glial cells into functional neurons around an implanted device. In this way, the ideal outcome would be to steer astrocyte differentiation away from a reactive, neurotoxic phenotype toward a potentially neuro-restorative and functionally integrated component of the local neural circuitry that ultimately improves device longevity and performance.

## Conclusion

Recent departures from traditional dogma that glial cells, including astrocytes, are not just passive support cells but can, in fact, actively regulate brain function down to individual synapses as well as between wider neural circuits have led to an elevated interest in developing innovative strategies aimed at modulating their behavior during CNS health and disease [24, 214]. However, previously established strategies directly targeting glial activation and scarring are currently being reassessed in light of new discoveries regarding the functional roles that astrocytes facilitate during CNS injury and repair. For instance, despite the widely understood idea that generation of a high-impedance glial scar is detrimental to recording and stimulating performance of intracortical microelectrodes [99], simply ablating reactive astrocytes before, during, or after scar formation may actually be detrimental to tissue recovery following CNS injury [132]. Indeed, positive correlations between both NeuN+ expression and GFAP+ immunoreactivity with recorded SNR ratios of chronically implanted devices suggest a more complex dynamic between astrocyte activation and neuronal firing activity following brain injury than previously recognized [96, 215]. Therefore, a deeper understanding of the regulatory mechanisms of astrocytes on modulating neuronal circuit function within the healthy and injured brain is required to fully realize their functional roles during wound healing and repair. This knowledge will, in turn, lead to more sophisticated device design and biomaterials approaches to modulating astrocyte behavior and improve the overall efficacy and performance of chronically implanted neural electrode interfaces.

This study investigated real-time spatiotemporal dynamics of astrocytes and vasculature in response to the intracortical microelectrode implantation. Astrocytes extend processes toward the implant as early as 1 h post-insertion. Following initial process extension, astrocytes cell bodies moved toward the probe around 12 h post-insertion. New astrocytes were observed within 24 h after probe implantation. Together, this led to astrocytic coverage of probe which increased by 4 d compared to 1 h post-insertion. In addition, astrocytes were less ramified and became hypertrophic, especially in the first 48 h. Importantly, soma area of astrocytes was correlated with the diameters of vessels suggesting a relationship to be further investigated in the future. These results highlight that astrocyte activity can be dynamic and multi-dimensional following probe implantation. Future investigations should focus on understanding the diverse roles and heterogeneity of astrocytes after device implantation in order to develop more sophisticated intervention strategies aimed at specific astrocyte responses. These efforts would benefit neuronal tissue outcomes after implantation and improve the device-tissue interface, ultimately paving the way for future advances in neural interfacing technology.

## Acknowledgements

The authors would like to thank Olivia Coyne and Derek Bashe for critical review of the manuscript. This work was supported by NIH NINDS R01NS094396 and a diversity supplement to this parent grant as well as NIH R01NS105691, R01NS115707, R21NS108098, F99NS124186, and NSF CAREER CBET 1943906.

## Contributions

Sajishnu Savya: Conceptualization, Methodology, Validation, Preliminary Analysis, Data curation, Investigation, Writing – outline, Visualization, Project administration.

Fan Li: Investigation, Visualization, Writing – original draft

Stephanie Lam: Formal Analysis, Data curation, Visualization,

Steven Wellman: Methodology, Investigation, Writing – review & editing.

Kevin C. Stieger: Investigation, Writing – review & editing.

James Eles: Supervision, Methodology, Writing – review & editing.

Keying Chen: Writing – review & editing.

Takashi D.Y. Kozai: Conceptualization, Methodology, Resources, Writing – review & editing, Supervision, Funding acquisition

## Supplementary Figures

**Supplementary Fig. 1.**
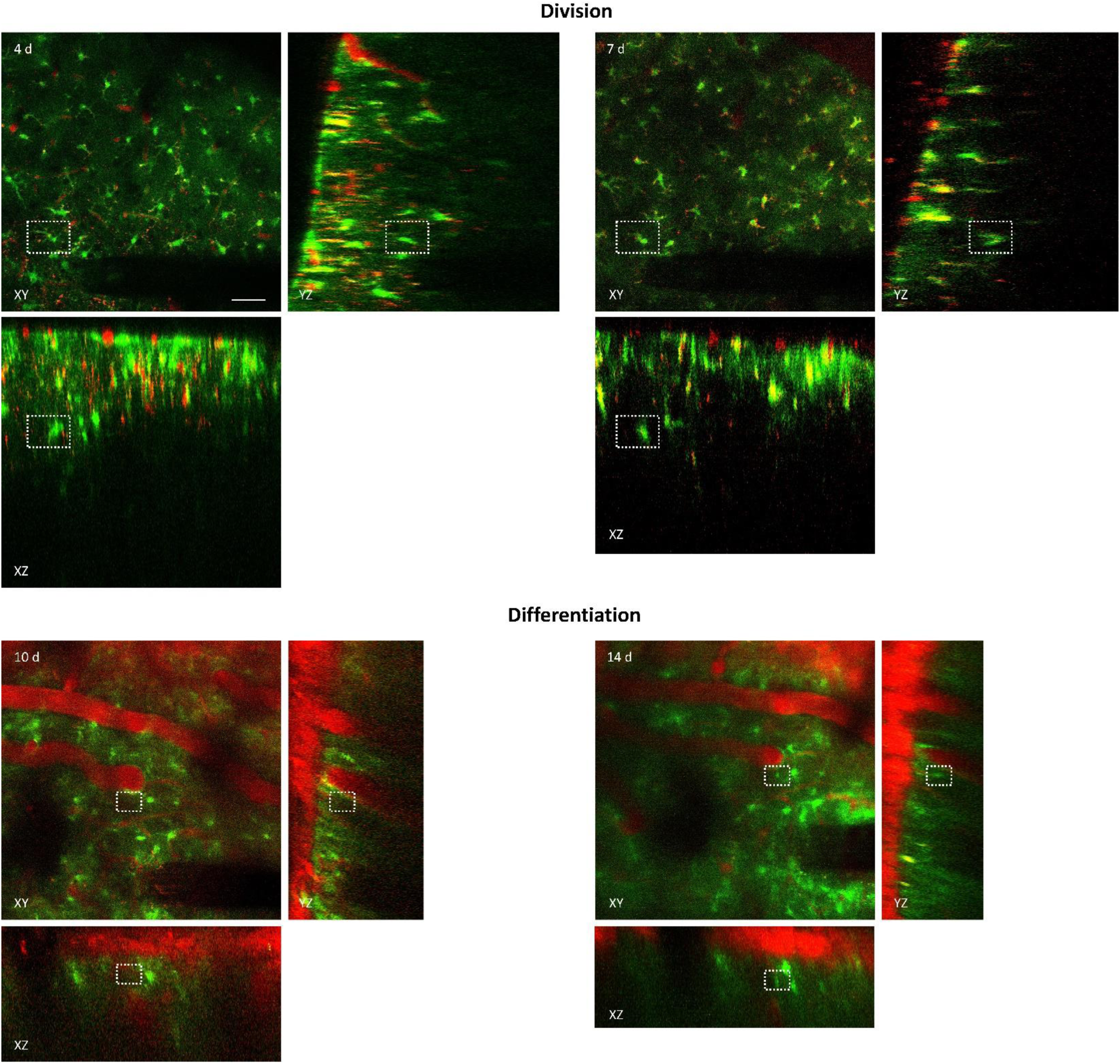
Validation of newly generated astrocytes in three-dimensional imaging volume. Top row: Orthogonal projection of astrocyte division in X-Y imaging plane around microelectrode at 4 and 7 d post-insertion. Two opposing astrocytes can be observed in all imaging planes at each imaging time point (*white rectangle*). Bottom row: Orthogonal projection of astrocyte differentiation event in X-Y imaging plane around microelectrode at 10 and 14 d post-insertion. A differentiated astrocyte can be observed as a gradual increase in fluorescence intensity within region of tissue previously lacking GFP+ signal (*white rectangle*). Note the absence of other astrocytes in surrounding tissue that could be mistakenly identified as differentiated cell. Scale bar = 50 μm.

**Supplementary Fig. 2.**
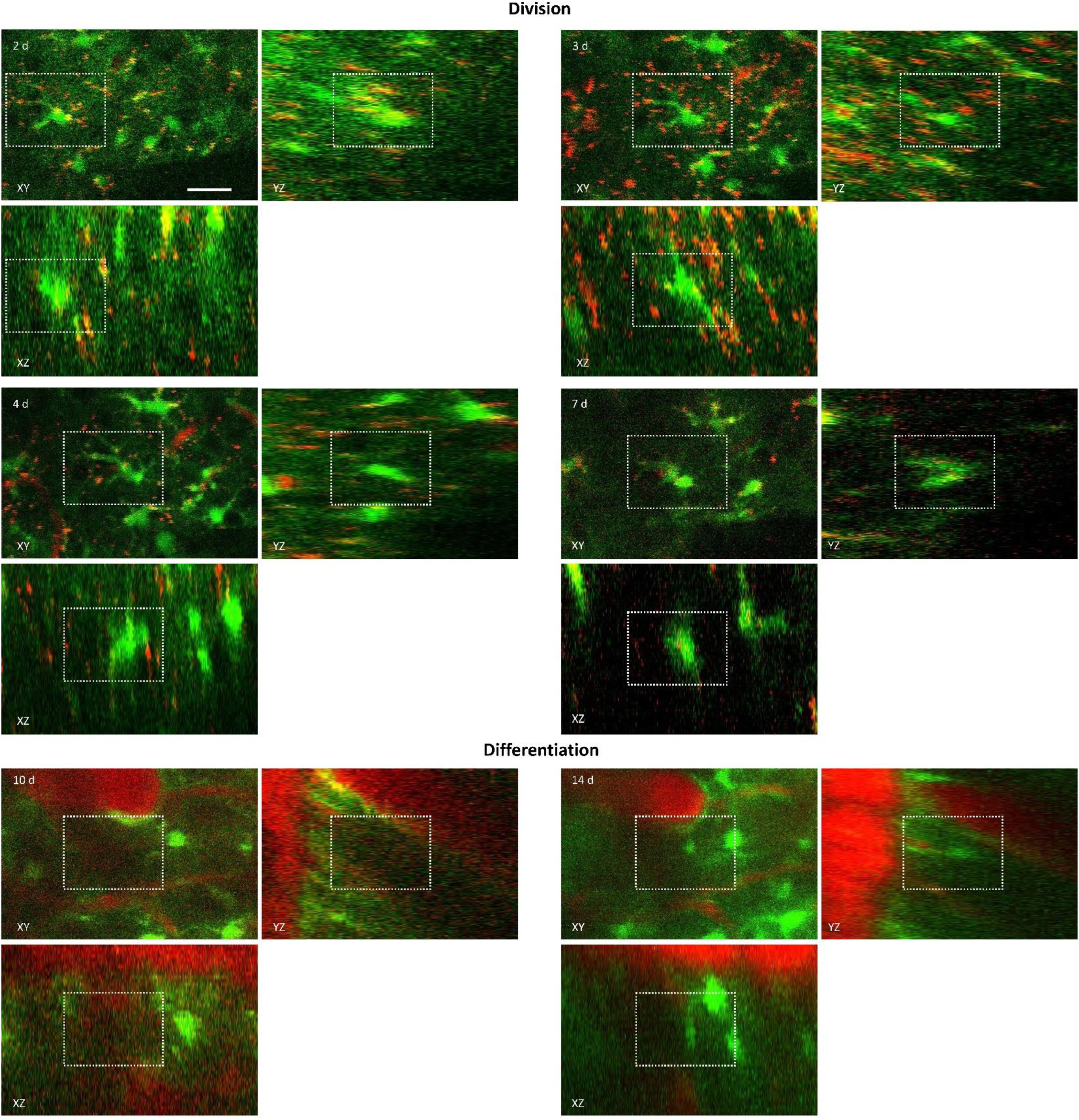
Enlarged view of division and differentiation in Supplementary Fig. 1. Top row: Orthogonal projection of astrocyte division in X-Y imaging plane around microelectrode at 2, 3, 4 and 7 d post-insertion. Two opposing astrocytes can be observed in all imaging planes at each imaging time point (*white rectangle*). Bottom row: Orthogonal projection of astrocyte differentiation event in X-Y imaging plane around microelectrode at 10 and 14 d post-insertion. A differentiated astrocyte can be observed as a gradual increase in fluorescence intensity within region of tissue previously lacking GFP+ signal (*white rectangle*). Note the absence of other astrocytes in surrounding tissue that could be mistakenly identified as differentiated cell. Scale bar = 25 μm.

**Supplementary Fig. 3.**
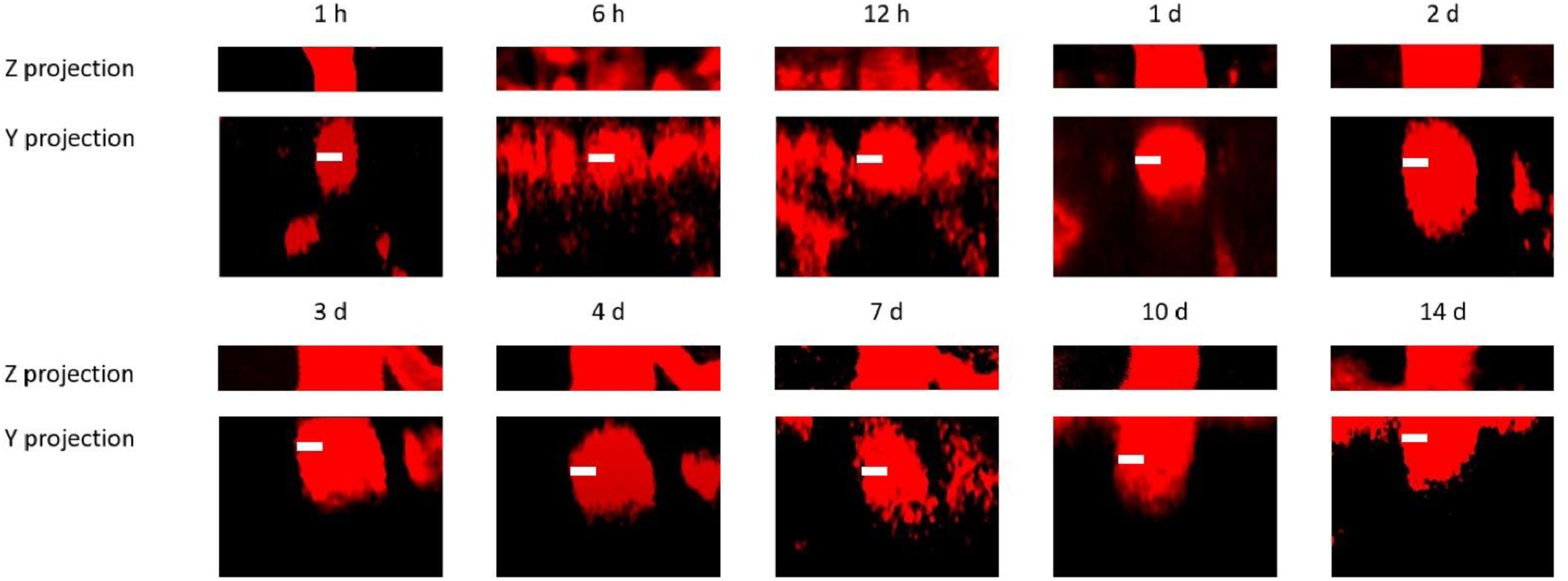
Validation of blood vessel diameter in Z projection and Y projection. Z and Y projection of blood vessel from 1 h to 14 d are shown. No significant drift is detected. Scale bar = 10 μm.

**Supplementary Fig. 4.**
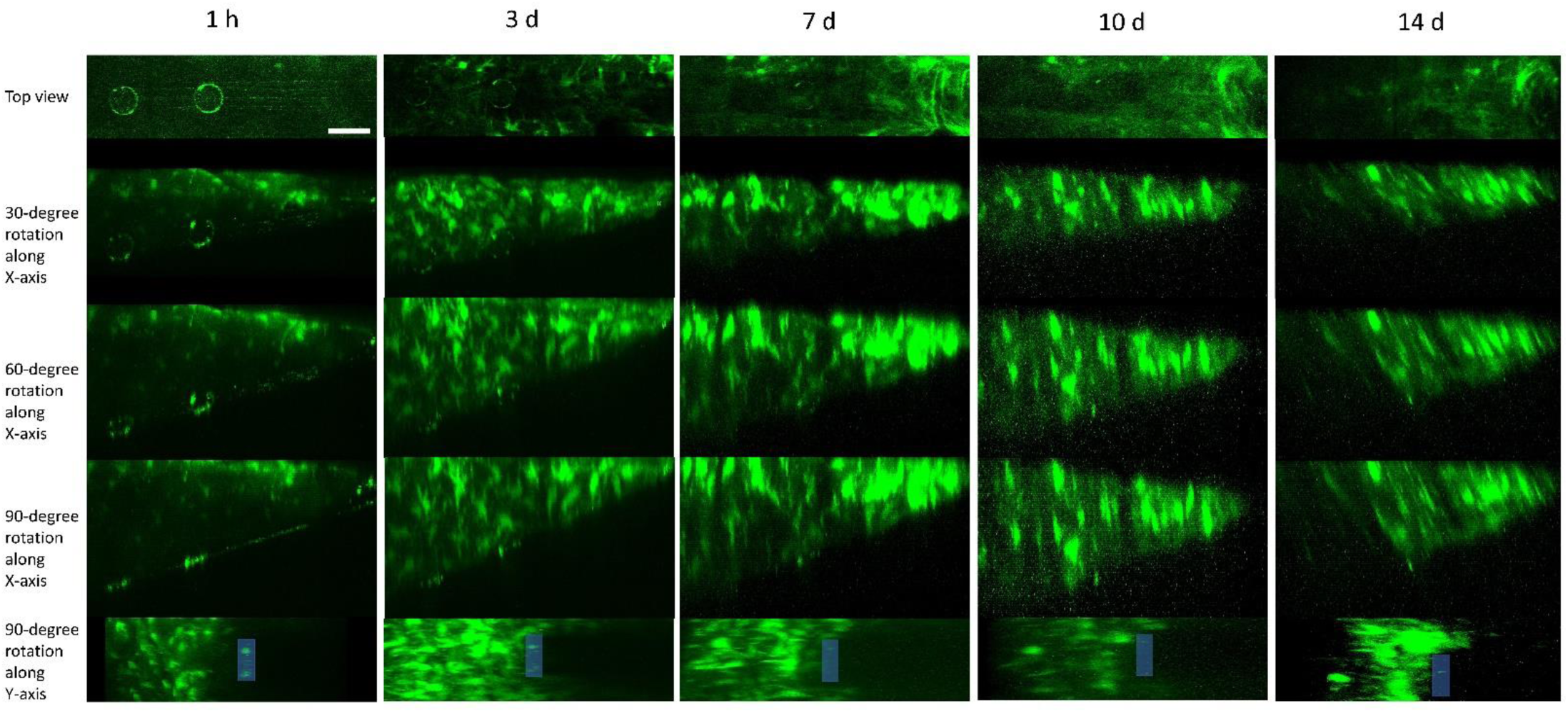
Validation of peri-implant cell population. Orthogonal projection of astrocyte population around microelectrode at 1 h, 3 d, 7 d, 10 d and 14 d are shown. Scale bar = 50 μm. Probe is outlined when in shaded blue.

## Footnotes

1 Reprinted from Biomaterial, Vol 164, Steven M.Wellman, Takashi D.Y.Kozai, *In vivo* spatiotemporal dynamics of NG2 glia activity caused by neural electrode implantation, Pages No. 121-133, Copyright (2018), with permission from Elsevier.

2 Reprinted from Biomaterial, Vol 164, Steven M.Wellman, Takashi D.Y.Kozai, *In vivo* spatiotemporal dynamics of NG2 glia activity caused by neural electrode implantation, Pages No. 121-133, Copyright (2018), with permission from Elsevier.

3 Reprinted from Biomaterial, Vol 164, Steven M.Wellman, Takashi D.Y.Kozai, *In vivo* spatiotemporal dynamics of NG2 glia activity caused by neural electrode implantation, Pages No. 121-133, Copyright (2018), with permission from Elsevier.

4 Reprinted from Biomaterial, Vol 164, Steven M.Wellman, Takashi D.Y.Kozai, *In vivo* spatiotemporal dynamics of NG2 glia activity caused by neural electrode implantation, Pages No. 121-133, Copyright (2018), with permission from Elsevier.

