## Supplemental Figures for "In vivo spatiotemporal dynamics of astrocyte reactivity following neural electrode implantation"

**Supplementary Figures**

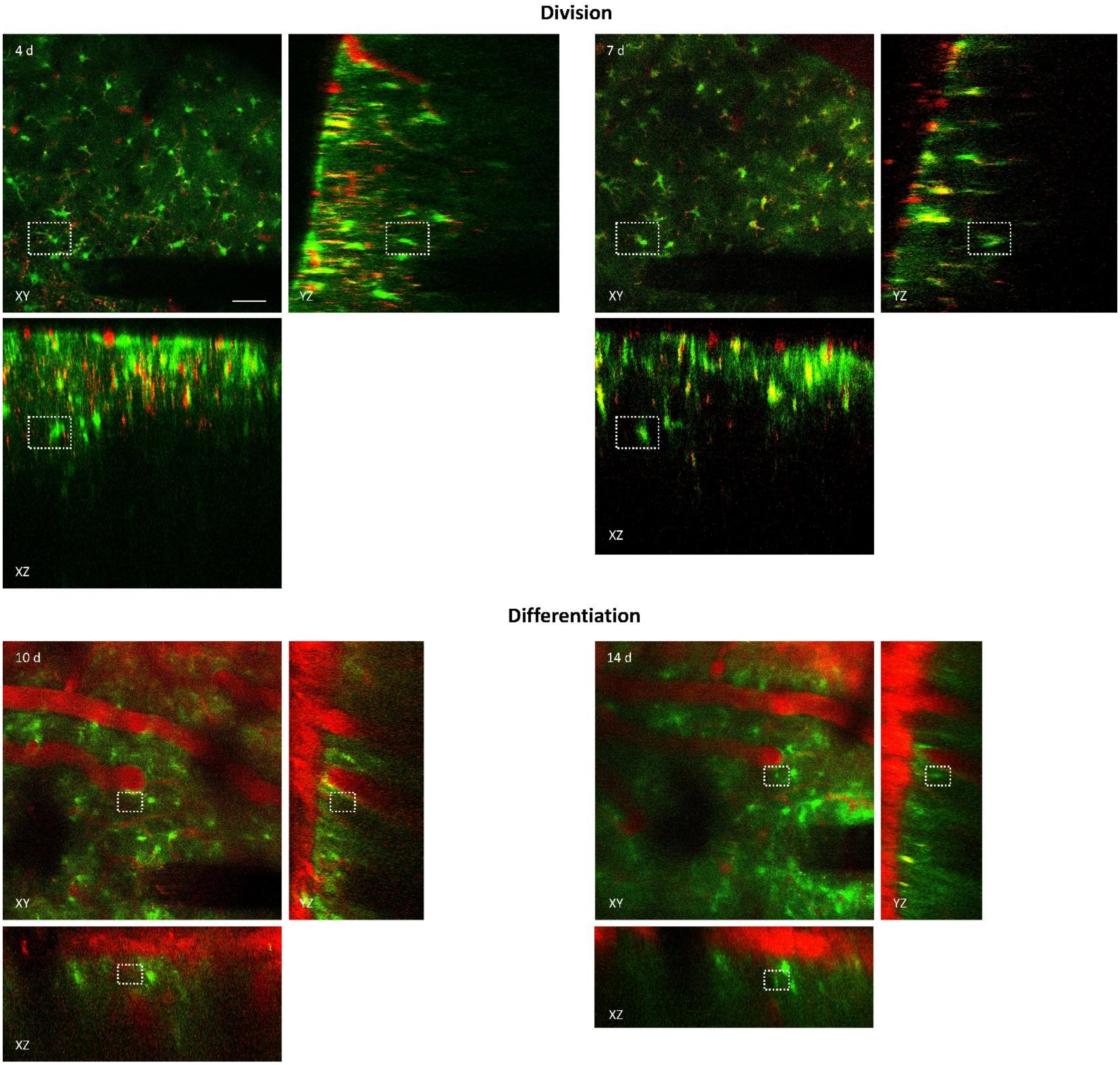


**Supplementary Fig. 1 Validation of newly generated astrocytes in three-dimensional imaging volume.** Top row: Orthogonal projection of astrocyte division in X-Y imaging plane around microelectrode at 4 and 7 d post-insertion. Two opposing astrocytes can be observed in all imaging planes at each imaging time point (*white rectangle*). Bottom row: Orthogonal projection of astrocyte differentiation event in X-Y imaging plane around microelectrode at 10 and 14 d post-insertion. A differentiated astrocyte can be observed as a gradual increase in fluorescence intensity within region of tissue previously lacking GFP+ signal (*white rectangle*). Note the absence of other astrocytes in surrounding tissue that could be mistakenly identified as differentiated cell. Scale bar = 50 μm.

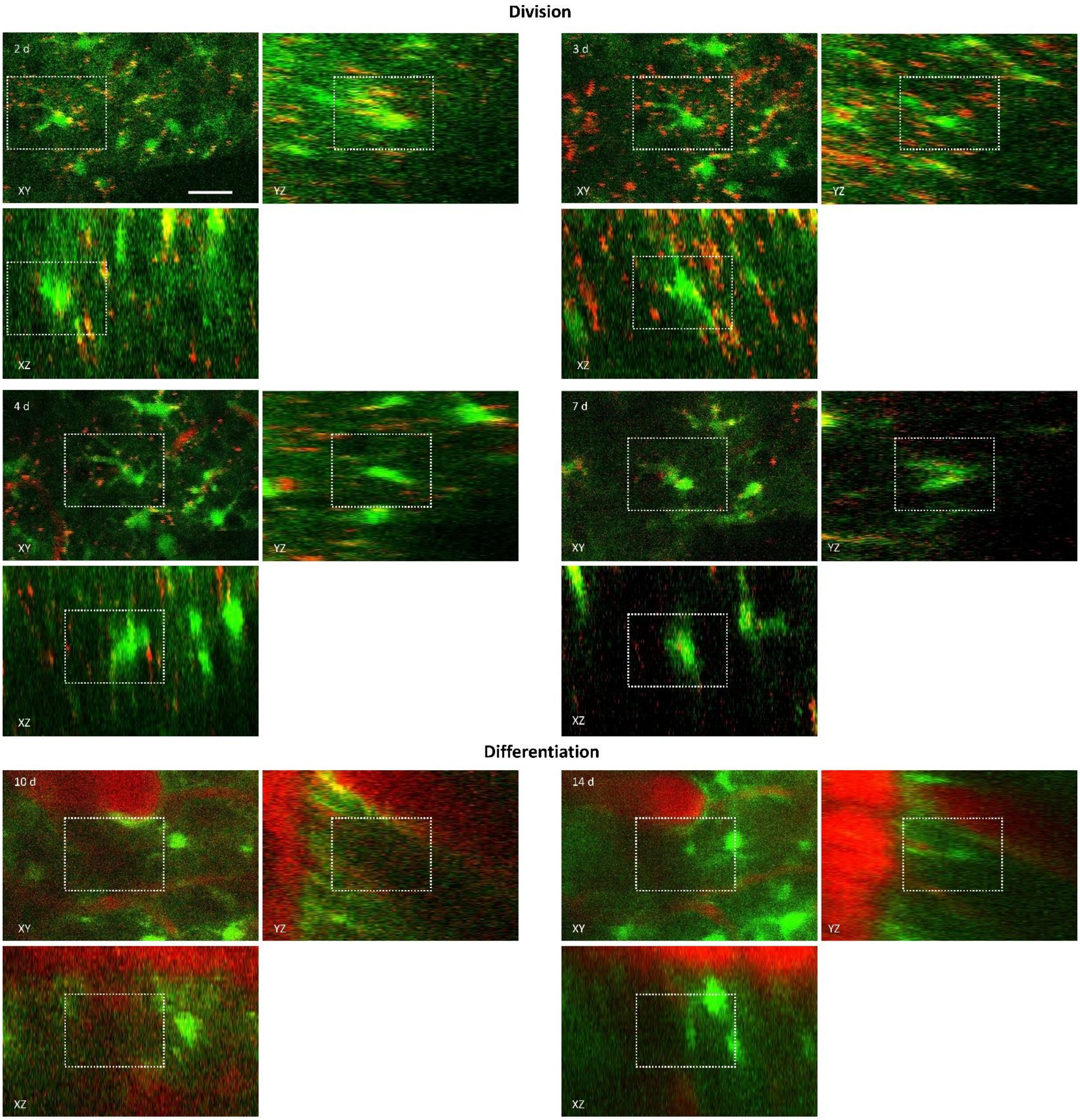


**Supplementary Fig. 2 Enlarged view of division and differentiation in Supplementary Fig. 1.**
Top row: Orthogonal projection of astrocyte division in X-Y imaging plane around microelectrode at 2, 3, 4 and 7 d post-insertion. Two opposing astrocytes can be observed in all imaging planes at each imaging time point (*white rectangle*). Bottom row: Orthogonal projection of astrocyte differentiation event in X-Y imaging plane around microelectrode at 10 and 14 d post-insertion. A differentiated astrocyte can be observed as a gradual increase in fluorescence intensity within region of tissue previously lacking GFP+ signal (*white rectangle*). Note the absence of other astrocytes in surrounding tissue that could be mistakenly identified as differentiated cell. Scale bar = 25 μm.

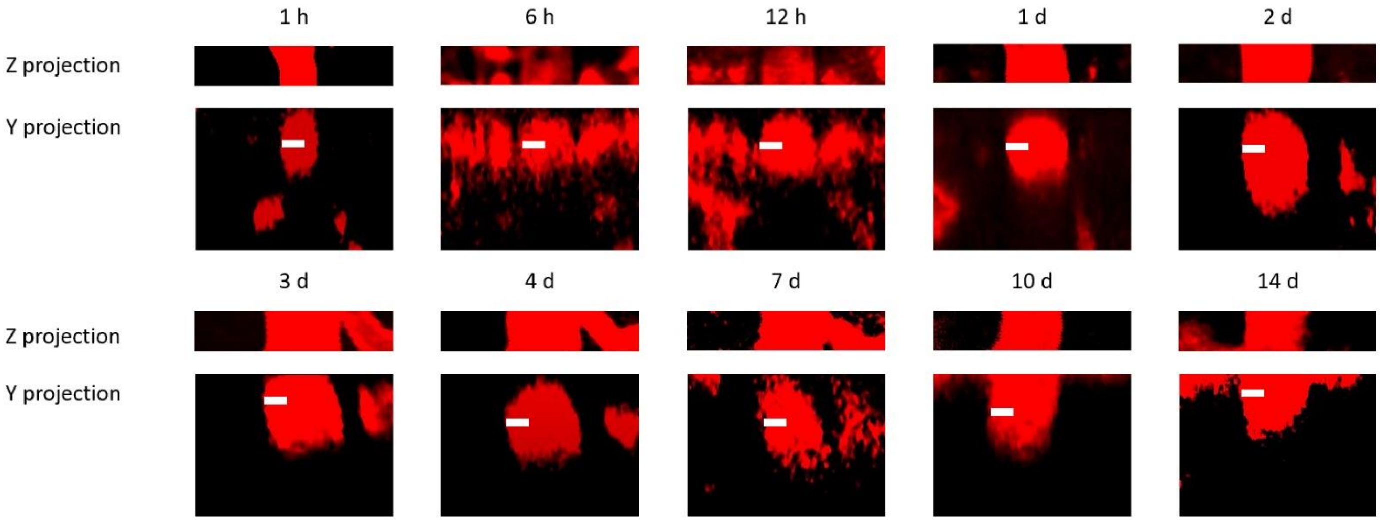


**Supplementary Fig. 3 Validation of blood vessel diameter in Z projection and Y projection.** Z and Y projection of blood vessel from 1 h to 14 d are shown. No significant drift is detected. Scale bar = 10 μm.

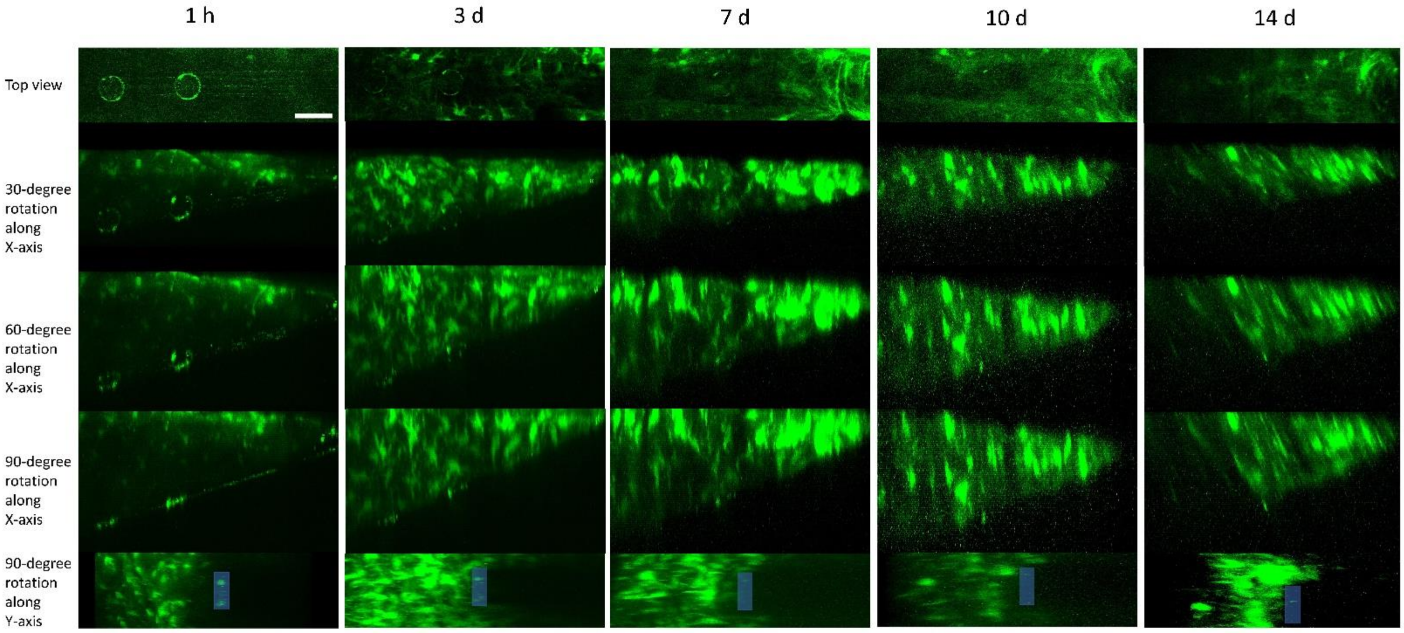


**Supplementary Fig. 4 Validation of peri-implant cell population.** Orthogonal projection of astrocyte population around microelectrode at 1 h, 3 d, 7 d, 10 d and 14 d are shown. Scale bar = 50 μm. Probe is outlined when in shaded blue.
